# Computational neurodevelopment: infant decision-making in changing environments

**DOI:** 10.1101/2024.10.08.617314

**Authors:** Rick A. Adams, Addison Billing, Levente Baljer, Eleanor Smith, Rob J. Cooper, Rebecca P. Lawson

## Abstract

In recognition of the fact that most psychiatric conditions have neurodevelopmental origins, there is an increasing interest in applying the methodological and conceptual approaches from computational psychiatry to developmental cohorts. However, the challenge of acquiring and modelling behavioural responses in very young infants has thus far proven difficult to overcome. To address this we developed a novel gaze-contingent, cued-reversal paradigm that allowed 6-10 month old infants to make overt behavioural responses to assess learning of expectations and updating of behaviour in response to change. We then fit computational models to infant behaviour and, for the first time, were able to validate the winning model to the same standards as would be expected of adults (e.g. good parameter recoverability, model identifiability and simulated behavioural responses). Similar to prior findings in adults, model-based prediction error measures correlated with post-switch increases in pupil size; consistent with noradrenaline’s hypothesised role in learning about change. Data-driven clustering based on model parameters revealed two infant behavioural subtypes hidden within the data; one with a perseverating profile and the other with a more exploratory decision-making pattern. This approach sheds new light on the ‘classic’ finding that all infants under 12 months tend to perseverate. Crucially, there were no significant differences in age between the clusters, but differences in terms of adaptive skills and temperament measured via gold-standard developmental assessments. These results prime the field for infant computational psychiatry, demonstrating that we can reliably fit models to infant data and that the parameters from such models can identify subgroups with distinct cognitive profiles that are superior to those derived from the behavioural data alone.

## Introduction

Our external environment is constantly changing and survival rests on the ability to recognise and respond to these changes. This basic cognitive capacity requires initial learning about statistical regularities (e.g., that A follows B) to build expectations about the world. However, sometimes the context changes (the order reverses, and B follows A), causing the experience of ‘prediction errors’ in service of new learning^1,2^. This is the cornerstone of cognitive flexibility and fundamental to adaptive behaviour^3^. Good cognitive flexibility is associated with better resilience to threats and negative life events^4,5^, higher educational attainment^6^, better quality of life^7^ and better social skills^8^. Compromised cognitive flexibility exists in almost every major neuropsychiatric and developmental condition to date^3,9,10^. This highlights the susceptibility of cognitive flexibility to risk factors that lead to adverse outcomes, and the importance of studying the emergence of error-driven learning at an early age.

Computational psychiatry has made great strides in parameterising the mechanisms underlying cognitive flexibility. To date, computational modelling has revealed that the learning mechanisms underlying expectation formation and updating are differentially affected in major depressive disorder^11,12^, autism spectrum disorder^13,14^, anxiety ^15–17^ and psychosis^18–20^. In fact, altered computations of learning under uncertainty have been identified as an important transdiagnostic feature across almost all forms of psychopathology^10,21,22^. This fundamental cognitive process depends upon the integrity of the ascending monoamines, with strong evidence for the role of the noradrenaline (NA) system. Often measured by-proxy with pupillometry^23–25^, NA is implicated in the encoding of ‘prediction errors’ signalling that a context has changed ^26–28^. These computational signatures of learning are also sensitive to medications that act via noradrenergic mechanisms ^27,29–31^, opening up potential new treatment options.

However, by studying the ‘end-state’ - adults who have a current psychiatric diagnosis - we risk only ever understanding the *consequences* and not the causes of neuropsychiatric disorders. This limits opportunities for intervention and prevention. We know that poor outcomes in later life (i.e. mental, neural and physical health difficulties) are preceded by changes in infancy, crucial neurobiological antecedents that are set in motion long before difficulties begin to emerge^32^. However, in order to meaningfully predict likelihood of cognitive difficulties or mental health challenges, we need predictors that speak to what the brain *does*: the computations it performs in response to, and in support of, our interactions with the environment.

Human infants are equipped with the capacity to learn statistical regularities about their environment^33^, and this is readily observed from about 6 months of age^34,35^. This fundamental mechanism is phylogenetically preserved across non-human primates, rodents and birds^36^; suggesting experience-dependent plasticity acts as scaffold for higher cognitive functions. Such an ability has been suggested to support the transition from reflexes (e.g., rooting/suckling) to inference (e.g., anticipating milk in sight of a bottle)^37^. Classic paradigms – such as Piaget’s A-not-B task – would suggest that infants under 12 months old typically do not show cognitive flexibility, tending to perseverate on previous choices^38^. However, measures of cognitive flexibility in infants are relatively blunt instruments (in-lab structured play with binary outcomes: e.g., infant ‘passed’ or ‘did not pass’)^39^. These measures are often confounded by social interaction with the experimenter and lack the granularity to capture meaningful individual or mechanistic differences between infants. Neural responses similar to prediction errors appear as early as 6 months of age ^35,40^. However, the trial-by-trial dynamics of learning have been difficult to measure in infants this young. Notably, standard tasks preclude computational interrogation of behaviour, a nontrivial issue affecting all developmental neuroscience, since infants < 12 months cannot follow explicit task instructions, make overt responses (e.g., button presses), or pay attention for more than a few minutes. These limitations have made it difficult to reliably and robustly model infant responses.

To overcome these challenges, we developed a novel decision-making paradigm involving a contextual switch, namely a cued reversal paradigm to capture infant behaviour from 6–10 months of age. This paradigm allows infants to interact with a visual scene and make choices (e.g., right or left) to reveal outcomes (e.g., a cartoon reward). This requires infants to form expectations about the outcomes under each context and deploy the appropriate response to be rewarded. We acquire infants’ choices trial-by-trial using their viewing behaviour via gaze-contingent eye-tracking and concurrently measure pupil size as a non-invasive proxy for phasic noradrenergic responses^24,25,28^. Reversal paradigms are ideal for testing the emergence of cognitive flexibility in infants, as making correct responses requires the maintenance of information about the context, recognising the change in the context and deploying the correct responses.

We then model behaviour on this paradigm with a Markov decision process (MDP) formulation of active inference and estimate model parameters specific to each infant. MDP models express environmental dynamics using discrete states and time and describe the mapping from states to outcomes using probability distributions. Active inference is a Bayesian framework that expresses attention, perception, and action in terms of minimization of an information-theoretic quantity, namely variational free energy (VFE), where VFE is an upper bound on surprise (or negative log model evidence) ^41,42^. The model responses are driven by a function called expected free energy, which incorporates both instrumental incentives and uncertainty-resolving motivations about the latent variables and model parameters. Crucially, to set the standards for infant computational psychiatry on solid foundations, we held this model to the same standards as behavioural modelling in adults, including multi-stage model comparisons, and demonstration of parameter recoverability, identifiability and the ability to accurately recapitulate infant behavioural responses with simulations^43,44^.

Based on prior literature in adults showing that pupil size encodes surprise ^26–28^, we wanted to assess whether a similar noradrenergic mechanism drives new learning in early life, and therefore hypothesised that infants’ model-derived estimates of prediction errors would drive increases in pupil size after the reversal. We then sought to combine behavioural and data-driven modelling approaches by clustering, based on the estimated parameters for each infant, to assess for subgroups expressing distinct profiles of cognitive flexibility. Finally, to give an indication of whether subgroups might differ in terms of the earliest indicators of developmental risk, we acquired gold-standard assessments of adaptive functioning and temperament which could be explained by computationally distinct mechanisms.

## Methods

### Participants

We recruited infants aged 5 to 10 months old who were carried to term and did not have any known or suspected developmental condition or delay. The participants were gifted little toys and t-shirts, and parents were reimbursed for travel expenses. The infants who completed at least three trials following the reversal were included in the analysis, as it is more likely that their expectations were violated in this interval. We recruited 55 infants, and 41 infants were included in the final analysis. Full demographics of the sample can be seen in **Table 1**. The gender ratio of females in sample was 0.54. A chi-square test revealed that the proportion of females and males were not different *χ*^2^(1,41) = 0.219, *p* = 0.639. Out of 14 excluded infants, 10 were either unable to perform the tasks or did not meet the data inclusion criterion. A further four infants were excluded due to a bug in the script that affected a subset of infants (see the supplementary materials).

**Table 1.**
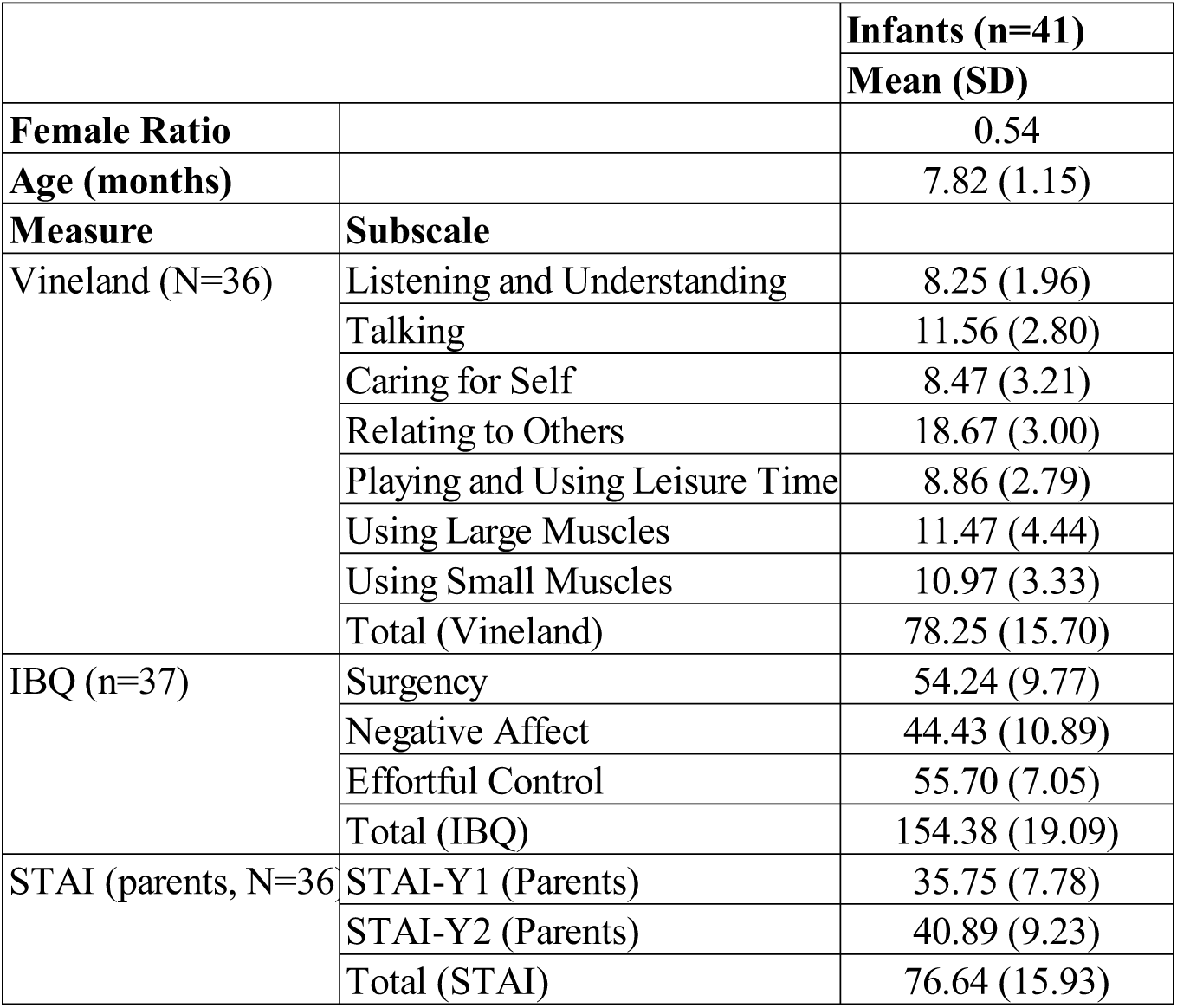
Means and standard deviation of the demographic data. Vineland: Vineland-II Adaptive Behaviour Scales, IBQ: Infant behaviour questionnaire, STAI: State-Trait Anxiety Inventory. Vineland and IBQ pertain to the infants, whereas STAI pertains to the parents. The number of participants who completed the questionnaires is reported separately for each questionnaire.

This study was approved by the University of Cambridge Psychology Research Ethics Committee (CPREC #PRE.2019.016). Participants were approached from the Cambridge BabyLab Infant Database, which recruits from flyers, advertisements and via linked studies at the local maternity hospital.

### Questionnaires

Adaptive functioning was assessed with the Vineland-II Adaptive Behaviour Scales and their temperament with the Infant Behaviour Questionnaire – Revised Very Short Forms (IBQ-R VSF). The Vineland-II questionnaire provides an measure of adaptive behaviour in communication, daily living, socialisation, and motor skills domains, and it has been widely used to assess the developmental state starting from birth and assist in the diagnosis of developmental impairments^45^. IBQ-R VSF is based on three dimensions of the IBQ-R encompassing the positive affect/surgency (PAS), negative affectivity (NEG) and Orienting/Regulatory Capacity (ORC) ^46^. We additionally collected the State-Trait Anxiety Inventory (STAI) scores on parents ^47^. Full details of questionnaire scores can be found in **Table 1**.

### Cued reversal paradigm

In this eye-tracking paradigm, the mapping between choices and outcomes changed abruptly during the experiment (i.e., reversal), forcing the decision-maker to adapt to the new context. Trials started with gaze-contingent orange and blue star cues which served as attention grabbers and triggered the start of each trial in a gaze-contingent manner. The cue colour indicated the phase in the paradigm (pre- or post-reversal). Upon looking at the coloured cues, they would spin for 500ms, followed by an auditory cue lasting 1000ms, and the decision-making period would begin 500ms after the onset of the auditory cue. The tone of the auditory cues were different for the pre- and post-reversal phases. The cue was followed by a 2000ms decision making screen in which two squares, namely the left and right regions of interest (ROIs), were displayed. One ROI was “correct” on a given trial and, if chosen, lead to a clear clip of an animation showing “five little monkeys”. Choosing the “incorrect” ROI lead to a blurred version of the same clip with the audio reversed. These served as rewarding and aversive outcomes respectively. The infants set the correct ROI in the pre-reversal phase by deploying a greater dwell time on one of the ROIs on the first trial. This meant that the infants’ decision on the first trial of each block was always correct. The correct and incorrect ROIs were swapped in the post-relative to the pre-reversal phase. The animation clip appeared for 5000ms at the ROI with the greatest dwell time, depending on whether a correct decision was made. At the end of each trial, there was a jittered baseline period between 5000-7000ms during which time concentric circles appeared and disappeared on screen and gentle sounds played. Each block consisted of eighteen trials, with nine pre- and post-reversal trials. There were two blocks of this paradigm, but not all infants completed the second block due to fussiness. During the task infants also wore a soft neoprene cap comprising hexagonal LUMO tiles for concurrent measurement of diffuse optical tomography (DOT) but those measurements are not reported here. Please see **Figure 1** for the experimental flowchart.

**Figure 1.**
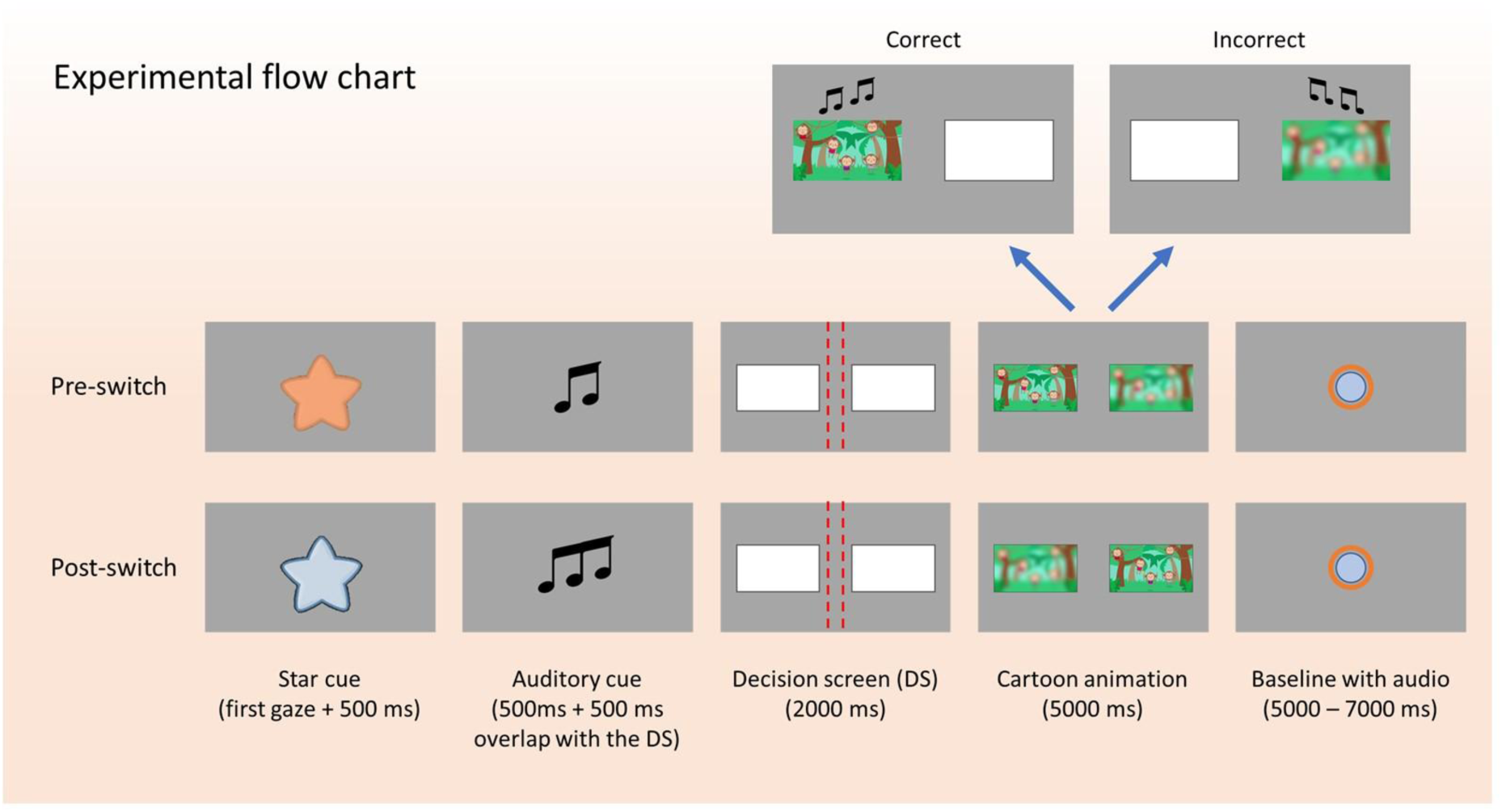
Cued reversal paradigm – The pre- and post-reversal trials started with gaze-contingent orange and blue star cues that span for 500ms upon looking at it. This was followed by a 1000ms long auditory cue that had a different tune in the pre- and post-reversal phases. The last 500ms of the auditory cue overlapped with the decision screen, which lasted 2000ms. In the decision screen, two white rectangles were displayed on either side of the screen. There were three ROIs, the left and right large ROIs and a narrow ROI at the centre (red dashed lines were not visible, and they are used for illustrative purposes only). The left and right large ROIs were defined as the area from the left red line to the left edge and the right red line to the right edge of the screen, respectively. The infants’ choices between the left and right ROIs were determined by dwell times, where the ROI with the greater dwell time was registered as the infants’ choice. The trials where the combined dwell time on the left and right ROIs was shorter than 500ms on the decision screen were repeated and no animation was played on these trials. In the cartoon animation period, a clear or blurred animation would play at the ROI of choice for 5000ms, depending on whether a correct decision was made. Correct and incorrect choices prompted the clear and blurred versions of the animation, and the audio was reversed in the blurred animation. Finally, there was a baseline period, which involved an animation of two concentric circles with an auditory tune, and its duration was jittered between 5000 – 7000ms. For illustrative purposes, we assumed that the left ROI was chosen on the first trial, thus setting the correct side of the screen as the left and right ROIs in the pre- and post-reversal phases, respectively.

### Eye tracking and pupillometry

The eye movements and pupillary responses of the infants were collected with an Eyelink 1000 plus eye tracker that used a 940 nm illuminator where a 17’’ monitor was mounted on a flexible arm, making it suitable for infant research. The sampling rate of the eye tracker was 500 Hz. We used the SHINE Toolbox to match the frames of the cartoon clip and the star cues in terms of luminance ^48^.

Previous work in the literature suggests that pupil dilation reflects surprise. Here we used pupillary responses as a proxy for surprise and tested whether there were any correlations between model-derived prediction errors and pupillary responses. We were interested in the pupillary responses from the animation period as the animations were used as feedback to decisions, and therefore, any violations in the infants’ expectations should manifest in this period. We first removed pupil data when the gaze was off-screen by replacing them with NaN entries and pre-processed the remaining data according to the steps outlined in Kret & Sjak-Shie (2019)^49^. Pupil data from the animation period on each trial was converted to z-scores using the mean and the standard deviation of the combined pupil data from the animation period of all trials for a given block. Average normalised pupil sizes over the animation period are reported across trials.

### Computational Modelling

Here, we briefly describe the states and outcomes in the MDP model, leaving the details to the **supplementary materials**. The MDP model of this paradigm consisted of a hierarchy with two levels. The higher level of the model involved a single state factor, an outcome modality labelled *context*, which encoded the pre- and post-reversal phases in the paradigm.

The lower level of the model was concerned with the dynamics on the within-trial level. There were three sets of states, namely *context*, *starting side* and *when*. The *context* states represented the pre- and post-reversal phases in the paradigm. The trials in the pre- and post-reversal phases always started with the orange and blue star cues, respectively, signalling the context. The *starting side* states described the infants’ choice between the left or right ROIs on the first trial of a block. The clear animations are displayed on the left and right ROIs in the pre-reversal phase, given that the infants choose the left and right ROIs on the first trial of each block, respectively. The ROI where the clear animation is displayed is swapped in the post-reversal phase. The *when* states represented the time and chosen actions in the decision-making period. These were labelled i) *t* = 1, ii) *t* = 2, *Left chosen*, and iii) *t* = 2, *Right chosen*. Time points *t* = 1 and *t* = 2 were associated with when the star cues (orange or blue) and animations (clear or blurred) were displayed on the screen, respectively. The states *Left chosen* and *Right chosen* described the chosen ROI in the decision period. There was a single outcome modality representing the *visual* features in the task, namely the orange and blue star cues and the clear and blurred versions of the animation.

We considered two kinds of generative models. These models included or excluded the *starting side* state factor. This state factor is necessary for the model to learn all unique combinations of state-to-outcome mappings (i.e., likelihood matrices). For example, consider an infant who chose the left ROI on the first trial of the first block. This means that the clear animation will play on the left ROI in the pre-reversal phase. If the same infant chooses the right ROI on the first trial of the second block, the clear animation will play on the right ROI in the pre-reversal phase. Given that the orange star cue is always associated with the pre-reversal phase, the infants can learn that choosing the left ROI after seeing the orange cue will prompt the clear animation in the first block. If they now carry forward these beliefs to the second block and keep choosing the left ROI after seeing the orange star cue, they will instead see the blurred version of the animation since they chose the right ROI on the first trial of the second block. Including the *starting side* factor in the model allows separate likelihood mappings for blocks 1 and 2, given that the infants may choose opposite ROIs on the first trials across blocks. For an infant who chooses opposite ROIs on the first trials across blocks, excluding this factor would force the model to overwrite its learned experiences associated with the pre-reversal phase of the first block with the pre-reversal phase experiences of the second block (just as all models have to overwrite their pre-reversal learning in the post-reversal phases). We referred to the models including and excluding the starting side factor, as *complex* and *simple* models, respectively.

### Model parameters

The parameters that were estimated on an individual basis by fitting the model to the data involved i) (inverse) volatility *ω*, preferences *c*, learning rate *η*, and ‘dominant policy bias’ (for choosing the left or right ROI) ɛ. A very high (inverse) volatility *ω* equips the agent with the belief that the environment is very rigid and the switches between contexts are unlikely, whereas a low *ω* means that the contextual switches are to be expected frequently (see Figure S3A). The parameter *c* determined the model’s preferences for clear and blurred animations, much like a reward sensitivity. The clear and blurred animations were assigned utilities *c* − 5 and 5 − *c*, respectively. Although we expected the infants to prefer the clear animation (*c* > 5), the formulation of the prior preference vector allowed preferences for the blurred animation as well (*c* < 5); see Figure S3B. Learning rate *η* modulated the rate at which the model learned the mapping from the states to the outcomes. The agent learned the likelihood mappings faster with high learning rate (see Figure S3C). Finally, the predominantly chosen ROI was calculated for each infant and block individually, and the most and least frequently visited ROIs were assigned ɛ and 10 − ɛ, respectively. These values expressed a bias for the dominant policy (i.e. a primacy bias) and were converted to probabilities by applying a softmax function (Left bias: **E** = σ([ ɛ 10 − ɛ]), Right bias **E** = σ([ 10 − ɛ ɛ])); see Figure S3D.

### Model comparison

In the first stage of model comparison, we compared the *complex* model and three variations of the *simple* model in terms of free energy (evidence). The first variation of the *simple* model carries its learned likelihood mappings from the first block forward to the second block. We will refer to this model as *simple – keep experience*. In the second variation, the likelihood mappings at the beginning of the second block were reset, and the learned mappings in the first block were discarded. We refer to this model as *simple – discard*. The third variation could not learn unique mappings from context states to outcomes. This model overwrites its experiences in the pre-reversal phase with the post-reversal phase, and the likelihood matrices under each context state are always identical. We will refer to this model as *simple – no context*.

The model’s choice behaviour is driven by a function called expected free energy. Expected free energy has three components, namely *novelty*, *epistemic value* and *extrinsic value*. In the second stage of model comparison, we fit models by excluding each component of the expected free energy – one at a time – in order to establish whether infants’ behaviour was motivated by each of these components. *Novelty* encourages the model to visit rarely occupied states to learn their mapping to the outcomes. A model without *novelty* would have no motive to resolve uncertainty about the mapping from states to the outcomes. *Epistemic value* encourages the model to sample the ROIs that would resolve uncertainty about the context. Excluding *epistemic value* robs the model of its drive to resolve uncertainty about the states. *Extrinsic value* drives the behaviour to fulfil the model’s preferences. A model without *extrinsic value* would have no preferences for the animations.

The final stage of model comparison involved estimating evidence for models defined in terms of parameter combinations. This approach employs a general linear model consisting of a term encoding group averages per parameter in a hierarchical inversion scheme, namely parametric empirical Bayes (PEB). The evidence for models including and excluding parameters is estimated using Bayesian model reduction (BMR)^50^. This method allows the posterior probabilities of the model parameters to be estimated. The parameters with low posterior probabilities (< 0.5) were fixed to their posterior expectations at the group level, and the model inversion was repeated for each infant.

In brief, model comparison involved three stages where different models were compared in terms of their free energy. In the first stage, we compared four models: i) *simple – keep experience*, ii) *simple – discard*, iii) *simple – no context*, and iv) *complex*. In the second stage, the winning model of the first stage was refitted to the data, excluding each component of the expected free energy (EFE) and compared with the original EFE. These models were i) *EFE*, ii) *EFE – without novelty*, iii) *EFE – without epistemic value*, and iv) *EFE – without extrinsic value*. In the third stage, models defined in terms of the combination of model parameters were evaluated using PEB and BMR. Because there were four free parameters, each of which can be either included or excluded to define a model, there were 2^4^ = 16 models to compare at this stage. The redundant parameters were fixed at their group-level means, and model inversion was repeated.

### Prediction errors

Prediction errors express a mismatch between the model’s expectations and its observations. Here, we define prediction errors on both levels of the model. Considering that the prediction errors from different levels of the model may contribute varying amounts to the infant’s physiological response after the reversal, we fit separate multilinear regression models involving the intercept term and a separate weight for the prediction errors to the normalised pupil data of each infant.

### Clustering

We applied k-means clustering on the estimated parameters to test if the clustering solution finds infant subgroups that are distinct in terms of their behavioural responses and questionnaire scores. Each parameter was rescaled using minmax normalisation over infants, and cosine distance was used to evaluate cluster membership. We tested clustering solutions that allowed up to 10 clusters in terms of the Calinski-Harabasz variance ratio criterion (VRC)^51^, where a greater VRC suggests a better clustering solution.

### Hypothesis testing

The behavioural responses in the cued reversal paradigm involved accuracy, perseveration, correct and incorrect dwell times (and their difference) in the decision period and the dwell time on the displayed animation. Accuracy is the binary correct or incorrect response on a given trial, where choosing the ROI with the clear animation is assumed to be the correct response. Perseveration is defined as incorrect responses on consecutive trials under the same context. For example, if the infants choose the incorrect ROI on two consecutive trials in the pre-reversal phase, that counts as perseveration. Dwell times correspond to the time spent looking at ROIs in the decision and animation periods. The left and right large ROIs were used to determine the dwell times. We used generalised linear mixed-effects models (GLMMs) for hypothesis testing involving the response variables accuracy, perseveration, dwell time – decision period and dwell time – animation period. See the **supplementary** for the details on GLMM analysis.

The differences between the pre- and post-reversal pupillary responses and prediction errors were tested using paired t-tests. The differences in questionnaire scores between clusters were evaluated using linear regression models that included cluster identity and age as regressors. Finally, the reported correlation coefficients correspond to the Pearson correlation coefficient.

## Results

### Behavioural responses

As expected, there was a main effect of reversal on accuracy *F*(1,1087) = 217.88, *p* < 0.001, and perseveration *F*(1,959) = 190.14, *p* < 0.001, with post hoc comparisons revealing that accuracy decreased and perseveration increased following the reversal (see **Figure 2A** and **B, Table 2**). Dwell time difference between the correct and incorrect ROIs in the decision period decreased *F*(1,81) = 87.94, *p* < 0.001 (see **Figure 2C, Table 2**) and the dwell time on the displayed animation also decreased in the post-compared to the pre-reversal phase *F*(1,51) = 29.26, *p* < 0.001 (See **Figure 2D, Table 2**). There was a main effect of block on perseveration *F*(1,959) = 8.55, *p* < 0.005 and a main effect of accuracy (correct vs incorrect) on the dwell time during the animation *F*(1,950) = 154.48, *p* < 0.001. Post hoc comparisons revealed increased perseveration in the second compared to the first block and increased animation dwell time on correct compared to the incorrect trials. See **Table 2** for full details of all post hoc comparisons.

**Figure 2.**
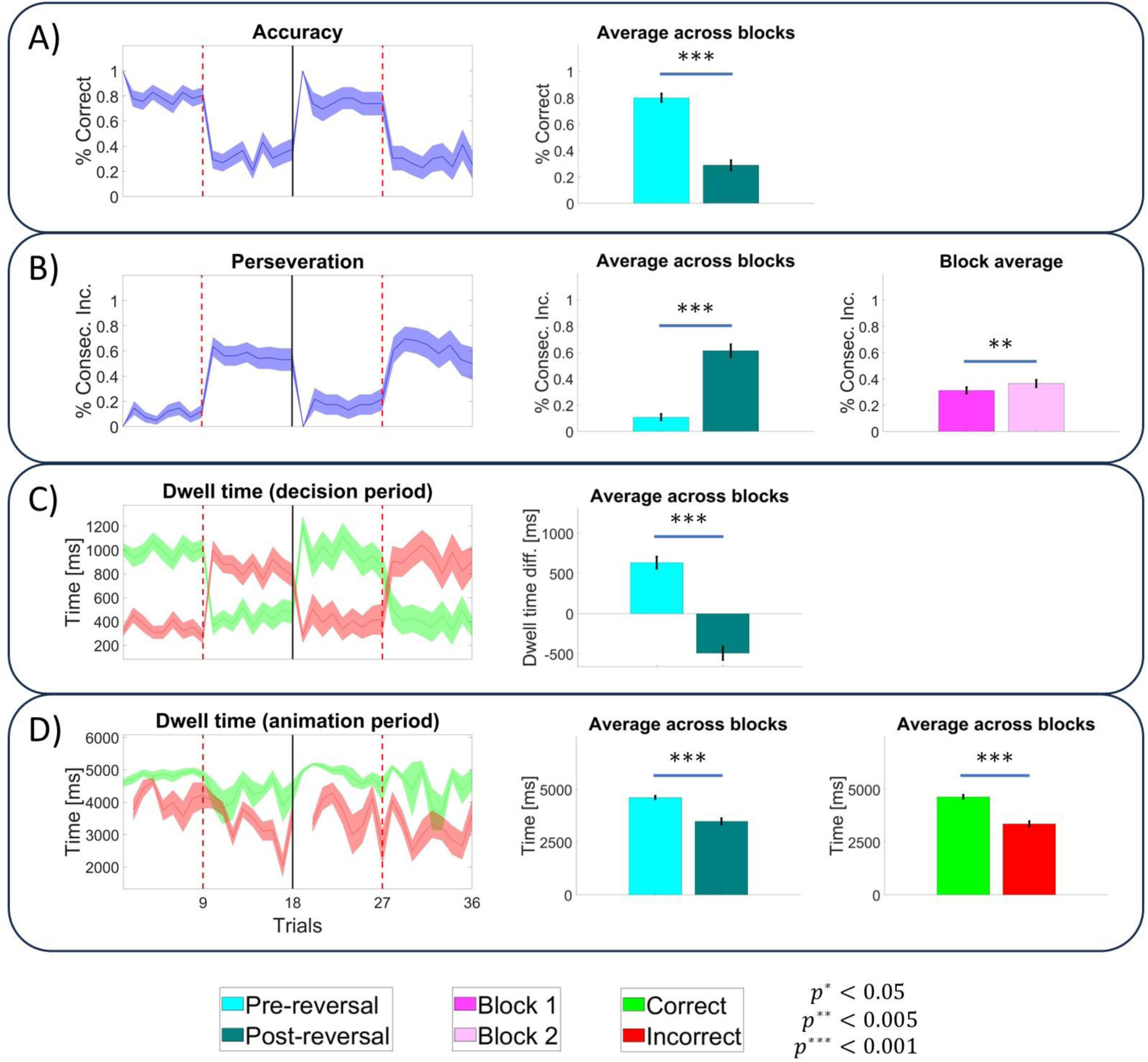
Behavioural measures – The panels on the left show behavioural measures across trials where the first and last 18 trials correspond to blocks 1 and 2, respectively. The red dashed lines indicate the final trial before the reversal in each block, while the solid black line at the 18^th^ trial indicates the end of the first block. The panels on the right illustrate the pairwise differences in behavioural measures with respect to different contrasts (see the legends at the bottom). A) The right panel shows average accuracy in the pre- and post-reversal phases. B) These panels show the average perseveration, defined in terms of consecutive incorrect responses in the same context (pre or post-reversal). The middle panel compares perseveration in pre-vs post-reversal phases averaged across blocks, whereas the right panel compares block-averaged perseveration (i.e., blocks 1 vs 2). C) The green and red shades in the left panel show the dwell time on correct and incorrect ROIs in the decision period, respectively. The right panel shows the dwell time difference between the correct and incorrect ROIs in the pre- and post-reversal phases, averaged across blocks. D) The green and red shades in the left panel show the dwell time on the displayed animation following correct and incorrect responses, respectively. Clear and blurred versions of the animation are displayed following correct and incorrect responses, respectively. This means that green and red shades are associated with the clear and blurred animations. The middle panel shows pre-vs post-reversal dwell time on the displayed animation, averaged across blocks. The right panel shows the dwell time on the animation following correct (clear animation) and incorrect (blurred animation) responses.

**Table 2.** Behavioural responses – This table shows the significant pairwise differences between behavioural measures for the contrasts i) post vs pre-reversal, ii) block 1 vs 2, and iii) correct vs incorrect trials.

| **Pairwise Comparisons** | **Contrast** | **95% CI** |  | **p-value** |
| --- | --- | --- | --- | --- |
|  |  | **LL** | **UL** |  |
| Accuracy | *Post - Pre* | -0.543 | -0.434 | < 0.001 |
| Perseveration | *Post - Pre* | 0.426 | 0.549 | < 0.001 |
|  | *Block 2 - Block 1* | 0.036 | 0.196 | < 0.005 |
| Dwell time difference - decision period [ms] | *Post - Pre* | -1355.927 | -881.281 | < 0.001 |
| Dwell time on cartoon - Animation period [ms] | *Post - Pre* | -802.239 | -367.955 | < 0.001 |
|  | *Correct - Incorrect* | 825.02 | 1134.392 | < 0.001 |

### Computational modelling

In this section, we compared models over three stages and reported the winning model at the group level. The winning model of the first stage was *simple – discard* (see **Figure 3A**). This model does not learn separate likelihood matrices for different starting sides as the *complex* model does. If infants chose opposite sides on the first trial of each block, there need to be four likelihood matrices to learn the true state–outcome mappings (see **Figure S2A**). The winning model, however, does not learn separate likelihood matrices for different starting sides, and it has only two likelihood matrices, one per context (i.e., pre- and post-reversal) state (see the right panel of **Figure S2B**). Furthermore, this model does not carry forward the learned contingencies from the first to the second block. This means that the learned contingencies are discarded at the end of the first block. The winning model of the second stage excludes *novelty* from the expected free energy (see **Figure 3B**). The final stage of model comparison shows that the 6^th^ model, which excludes (inverse) volatility *ω* and learning rate *η*, has the greatest evidence (see the top right panel of **Figure 3C**). We fixed the volatility *ω* and learning rate *η* parameters to their posterior expectations at the group level and repeated the inversion for each infant. Note that this model still incorporates volatility and learns from its recent experiences, but the parameters *ω* and *η* do not express between-subject differences because they are fixed to their group-level means.

**Figure 3.**
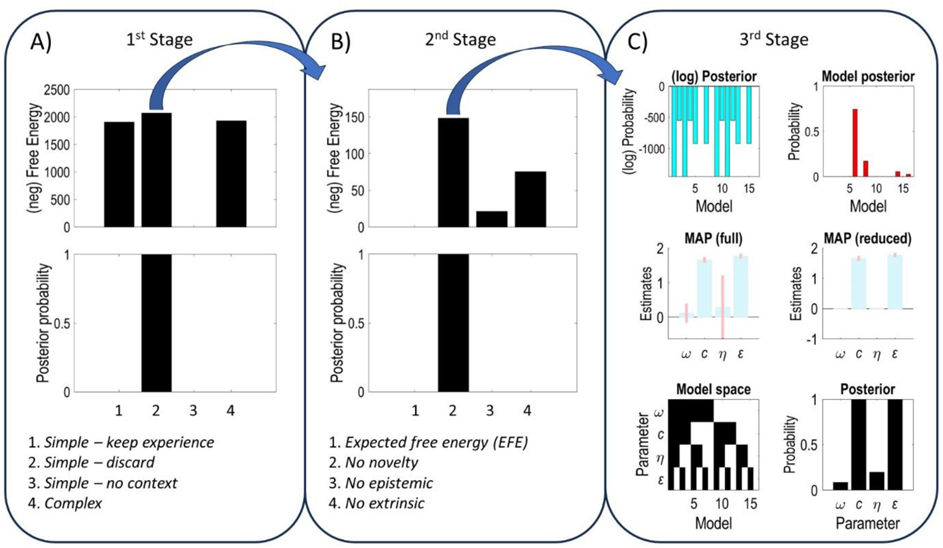
Model comparison – A) There are four competing models at this stage, namely i) simple – keep experience, ii) simple – discard, iii) simple – no context and iv) complex. The upper panel shows that *simple – discard* has more evidence than the other models, and the lower panel shows this model has a posterior probability of nearly 1. B) The winning model of the previous stage, *simple – discard*, is refitted to the data, excluding one component of expected free energy at a time. The competing models at this stage are i) expected free energy (EFE), ii) no novelty, iii) no epistemic and iv) no extrinsic. The *no novelty* model has more evidence than the rest and similarly has a posterior probability of nearly 1. C) The winning model of the second stage enters this stage as a parent model, where the children models are defined by turning off the parameters of the former. There are 2^4^ = 16 children models because each parameter of the parent model can be either included (on) or excluded (off). The bottom left panel shows the model space where the dark and white colours indicate which parameters are on or off for a given model. The top left and right panels compare models in terms of their evidence and posterior probabilities, respectively. The middle left and right panels show the maximum a posteriori (MAP) parameter estimates (in log space) at the group level before and after applying Bayesian model reduction, respectively. The posterior probabilities of parameters are presented at the bottom right panel. Parameters with higher posterior probabilities have greater evidence to be included in the model.

The left panel of **Figure 4A** shows the individual estimates of parameters *c* and ɛ for each infant (in log space). The right panel of **Figure 4A** shows a weak positive but statistically insignificant correlation (*r*(28) = 0.23, *p* = 0.15) between the parameters *c* and ɛ. The left and centre panels of **Figure 4B** show the group-level estimates of these two parameters and their posterior probabilities, respectively. We computed the degree of correspondence between the model’s and the infants’ choices by forcing the model to make the same observations and take the same actions as the infants in the first 1 to *n* − 1 trials and compared the model’s response to the infant’s decision on the *n*^*th*^ trial. We repeated this for each trial (except the first) and infant. The right panel of **Figure 4B** shows a median decision fit of ≈ 86%, with a minimum fit of 60%. To test the validity of the estimated parameters, we generated two datasets per infant using their individual parameter estimates and refitted the model to the simulated data. The original and the simulated datasets are presented in **Figure 4C**. There were high and significant correlations between the parameter estimates obtained from the first (original data) and the second (simulated data) inversions (see **Figure 4D**).

**Figure 4.**
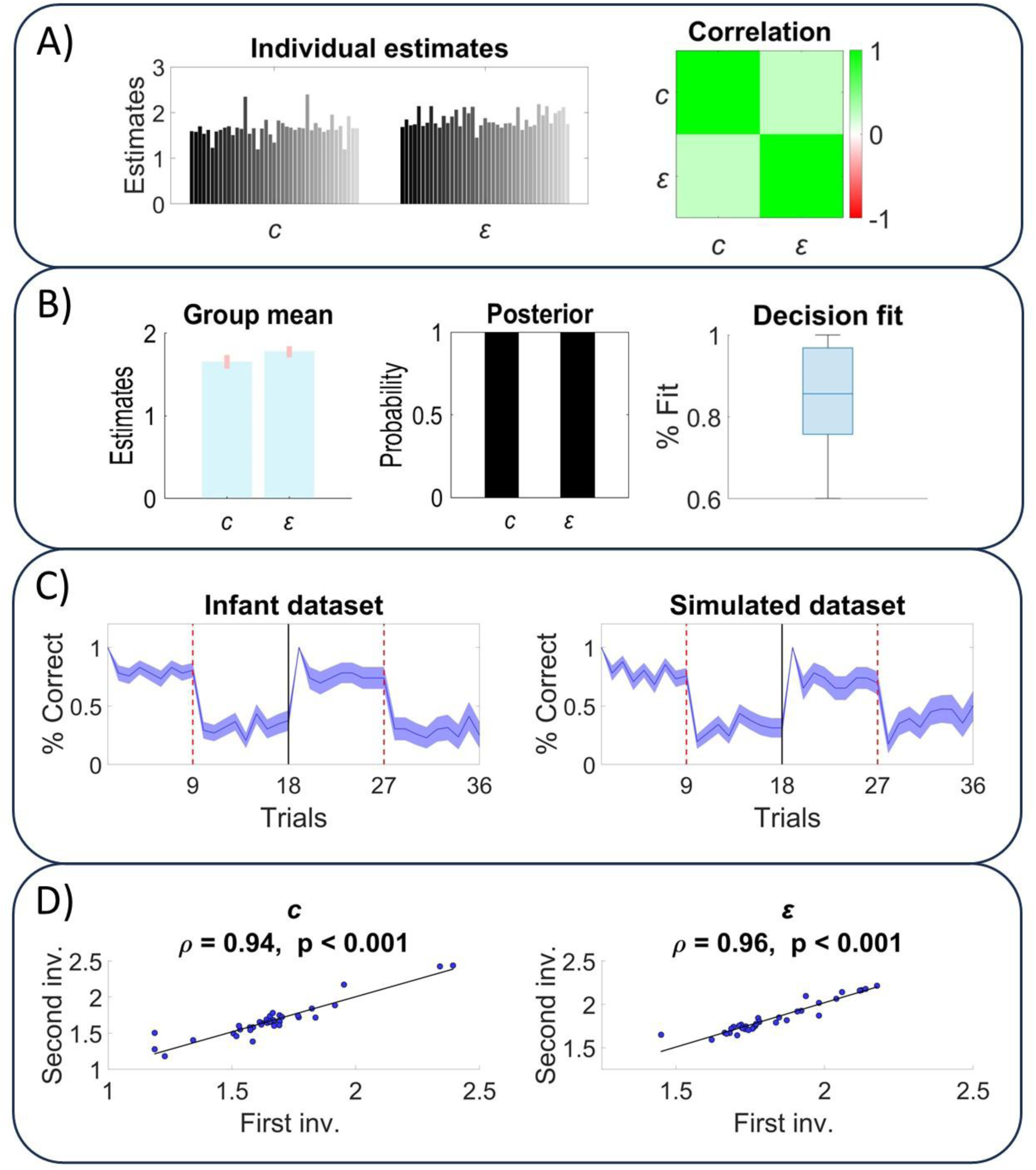
Parameter estimates and correlations – A) Individual parameter estimates and the correlation between the parameters are displayed on the left and right panels, respectively. B) The group-level means of parameters and their posterior probabilities are shown on the panels on the left and centre. The correspondence between the model’s and infants’ choices is presented in terms of decision fit on the right panel. C) Two datasets were generated per infant using their individual parameter estimates. The panel on the left shows the accuracy in the original dataset, whereas the right panel shows the average accuracy (over datasets and infants) in the simulated dataset. D) This panel presents the correlations between the original parameter estimates and the estimates obtained by fitting the model to the simulated dataset. Parameters are presented in log space.

### Pupillary responses and prediction errors

The left panels of **Figure 5A** show the normalised pupil sizes and prediction errors across trials. The prediction errors shown in the figure are a weighted linear combination of both high and low level prediction errors from the regression model. The pupil sizes and prediction errors increase in the few trials following the reversal. The pupil size profile in the second block shows a delayed increase and more variance – potentially due to the decreased number of infants in this block. The right panels show the difference between the average responses in the post and pre-reversal phases as a function of the number of trials *n* used to compute the averages in the respective phases. In the pre- and post-reversal phases, *n* preceding and following trials are used from the switch trial to compute the averages, where *n* ∈ {1,2, …, 9}. The average pupil size and prediction error follow a similar profile such that there is a substantial increase for *n* = 2, and they gradually decrease with increasing *n*. However, carrying out separate paired t-tests to test for the effect of reversal on average pupil sizes for *n* ∈ {1,2, …, 9}, *n* = 3 maximised the t-statistic.

**Figure 5.**
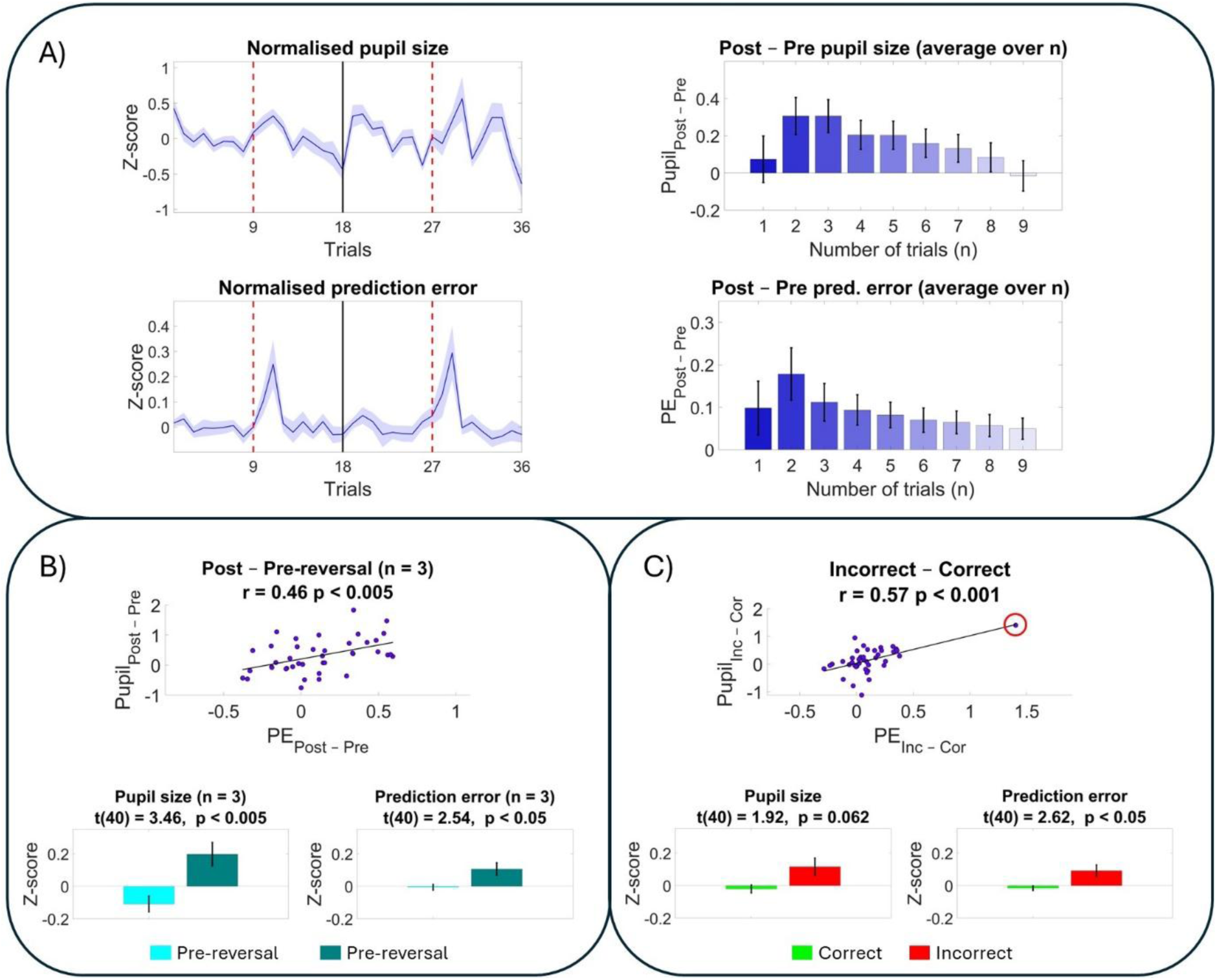
Pupil size and prediction error – A) Normalised pupil size and prediction errors are plotted across trials on the left panels. The differences between the average post and pre-reversal responses are plotted as a function of the number of trials (*n*) used before (pre-reversal) and after (post-reversal) the contextual switch. The average responses in the pre-reversal phase are obtained by computing the mean responses over *n* preceding trials relative to the switch trial. Similarly, *n* trials are used from the switch trial to compute the average responses in the post-reversal phase. The differences between the two are reported for the pupil size *Pupil*_*Post*−*Pre*_ and prediction errors *PE*_*Post*−*Pre*_ as a function of *n* on the right panels. B) The top panel shows that there is a moderate correlation between *Pupil*_*Post*−*Pre*_ and *PE*_*Post*−*Pre*_ for *n* = 3. The bottom panels show that there are significant differences between the post and pre-reversal pupil size and prediction errors for *n* = 3. C) The average difference between the responses on incorrect and correct trials are computed for pupil sizes *Pupil*_*Inc*−*Cor*_ and prediction errors *PE*_*Inc*−*Cor*_. The top panel shows a significant correlation between *Pupil*_*Inc*−*Cor*_ and *PE*_*Inc*−*Cor*_. Excluding the outlier infant (see the red circle), still yielded significant correlations. The bottom panels show that there is an increase in the pupil size and prediction errors on incorrect trials. However, this increase is only significant for the prediction error. The error bars show the standard error of the mean.

Carrying out paired t-tests using the average pupil sizes over *n* = 3 post (*M* = 0.20, *SD* = 0.46) and pre-reversal trials (*M* = −0.11, *SD* = 0.31) yielded a significant difference *t*(40) = 3.46, *p* < 0.005. Similarly, there was an increase in the prediction errors in the post (*M* = 0.11, *SD* = 0.25) compared to the pre-reversal trials (*M* = −0.01, *SD* = 0.11) when *n* = 3, *t*(40) = 2.54, *p* < 0.05. The differences between average responses in the post and pre-reversal phases are computed over *n* trials for pupil sizes *Pupil*_*Post*−*Pre*_ and prediction errors *PE*_*Post*−*Pre*_. There was a significant correlation between *Pupil*_*Post*−*Pre*_ and *PE*_*Post*−*Pre*_ for *n* = 3, *r*(39) = 0.46, *p* < 0.005. See **Figure 5B**.

Conducting paired t-tests using the average pupil sizes on incorrect (*M* = 0.12, *SD* = 0.34) and correct trials (*M* = −0.02, *SD* = 0.15) showed that there was a tendency for increased pupil size on incorrect trials *t*(40) = 1.92, *p* = 0.062. As one would expect, there was an increase in the size of prediction errors on incorrect (*M* = 0.09, *SD* = 0.23) compared to correct trials, (*M* = −0.02, *SD* = 0.09), *t*(40) = 2.62, *p* < 0.05. The differences between average responses in the incorrect and correct trials are calculated for pupil size *Pupil*_*Inc*−*Cor*_ and prediction errors *PE*_*Inc*−*Cor*_. There was a significant correlation between *Pupil*_*Inc*−*Cor*_ and *PE*_*Inc*−*Cor*_, *r*(39) = 0.57, *p* < 0.001. Excluding the outlier infant (see the red circle in **Figure 5C**) still yielded a significant correlation between *Pupil*_*Inc*−*Cor*_ and *PE*_*Inc*−*Cor*_ *r*(38) = 0.40, *p* < 0.05. See **Figure 5C**.

### Clustering

Applying k-means clustering on the minmax normalised parameter estimates yielded two infant clusters, shown with magenta (*N*_*Magenta*_ = 26) and yellow (*N*_*Yellow*_ = 15) colours in **Figure 6**. The top left panel of **Figure 6A** illustrates these two clusters in the parameter space *c* against ɛ. The boundary between these two clusters of infants is obtained by applying Fisher’s discriminant analysis^49^ (see the red and blue coloured areas). We included the main effect of cluster and the interaction between cluster and reversal in our GLMM analysis. This resulted in a significant interaction effect between cluster and reversal on the accuracy *F*(1,1085) = 46.45, *p* < 0.001, perseverance *F*(1,957) = 33.27, *p* < 0.001 and dwell time difference between the correct and incorrect ROIs in the decision period *F*(1,75) = 28.63, *p* < 0.001. Post hoc comparisons revealed increased accuracy and dwell time difference in the pre-reversal phase and decreased accuracy and dwell time difference in the post-reversal phase for the yellow compared to the magenta cluster. Post hoc comparisons also showed increased perseveration in the post-reversal phase for the yellow compared to the magenta cluster. See **Table 3A** and **Figure 6A**.

**Figure 6.**
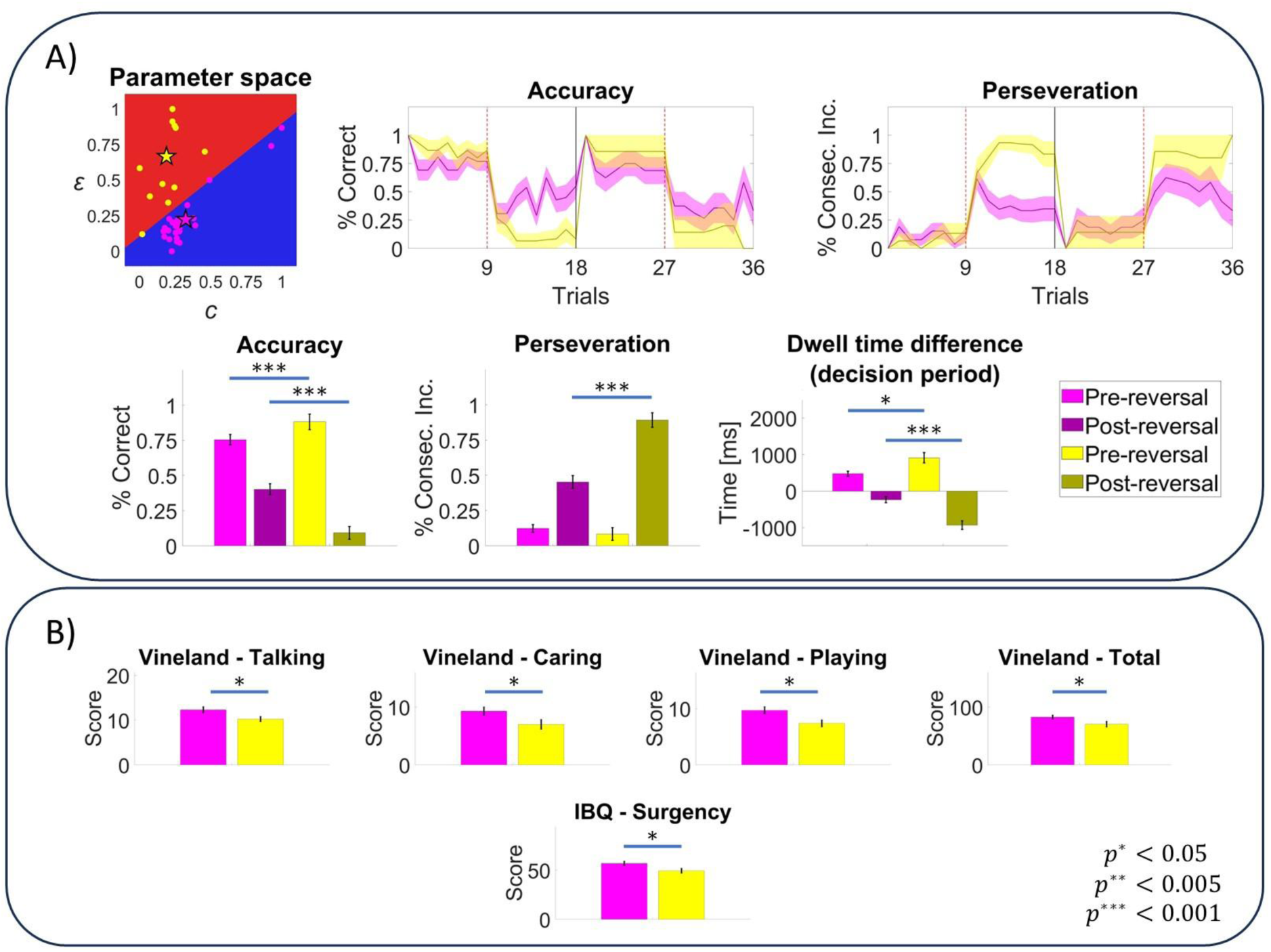
Infant clusters – A) Applying k-means clustering returned two clusters of infants, represented by the yellow and magenta colours. The top left panel shows the clusters in the parameter space after minmax normalisation, where the cluster centers (centroids) are shown with the star symbols. The differences between the two clusters manifest behaviourally in terms of accuracy, perseveration, and dwell time difference between the correct and incorrect ROIs in the decision period. The lower panels compare the clusters in the pre-reversal (light colours) and post-reversal (dark colours) phases. B) These panels show cluster-wise differences in terms of questionnaire scores. These are the Vineland subscales – Talking, – Caring, – Playing and the total Vineland score, and the IBQ – Surgency subscale from the Infant Behaviour Questionnaire (IBQ).

**Table 3.**
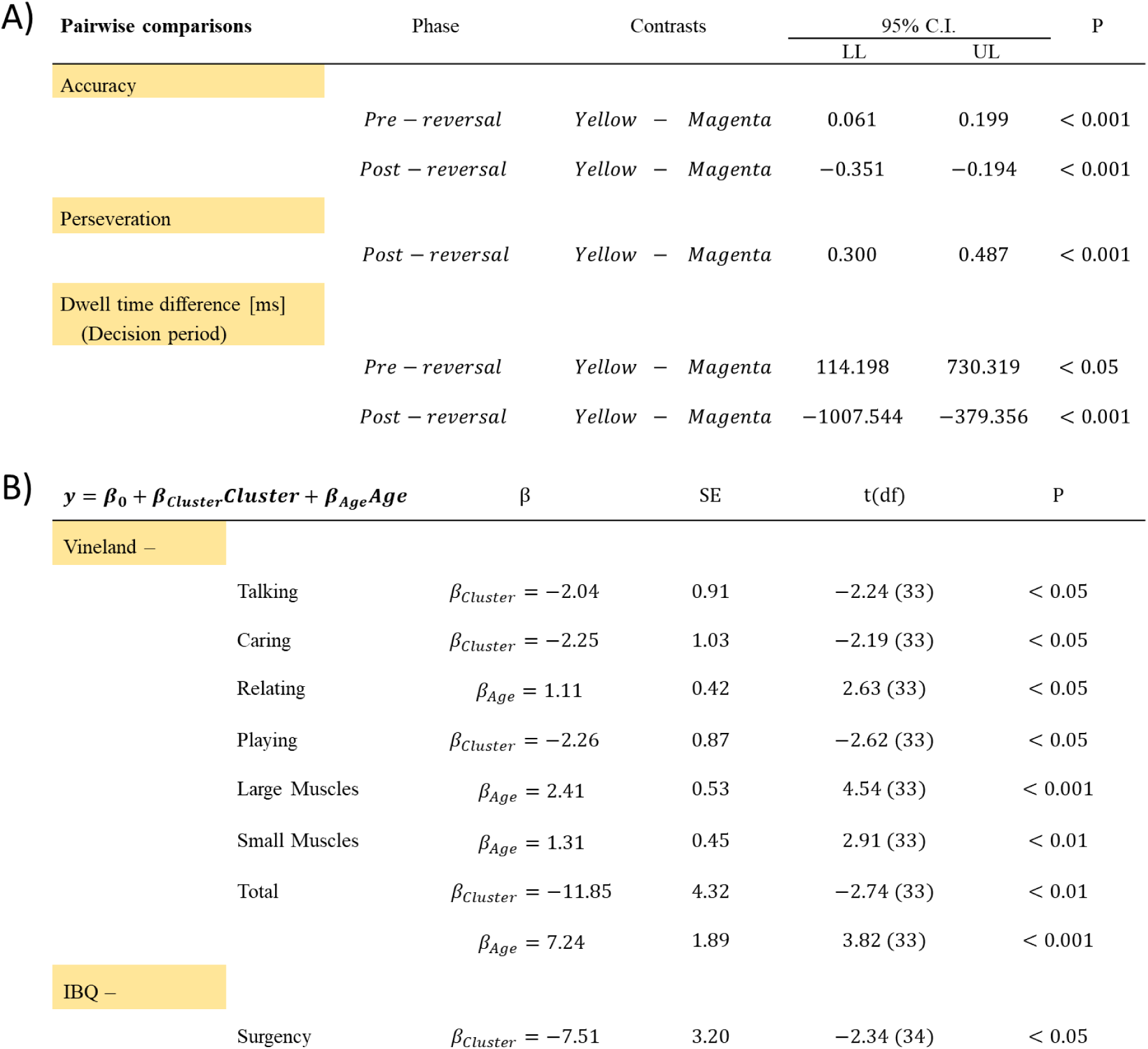
Cluster-wise differences – A) GLMM analysis of behavioural measures returned a significant interaction effect between the phase (pre- or post-reversal) and cluster identity on accuracy, perseveration and dwell time difference (decision period). This table shows the significant pairwise differences between clusters in the pre- and post-reversal phases. B) Multiple linear regression was used to test if the cluster identity and age explained questionnaire scores. The significant coefficients for cluster identity and age are reported in this table. The reference cluster is magenta for the cluster coefficient *β*_*Cluster*_, and negative coefficient estimates indicate reduced scores for the yellow cluster. Age is converted to months for the regression analysis.

| A) Pairwise comparisons | Phase | Contrasts | 95% C.I. |  | P |
| --- | --- | --- | --- | --- | --- |
|  |  |  | LL | UL |  |
| Accuracy | *Pre – reversal* | *Yellow – Magenta* | 0.061 | 0.199 | < 0.001 |
|  | *Post – reversal* | *Yellow – Magenta* | −0.351 | −0.194 | < 0.001 |
| Perseveration | *Post – reversal* | *Yellow – Magenta* | 0.300 | 0.487 | < 0.001 |
| Dwell time difference [ms]<br>(Decision period) | *Pre – reversal* | *Yellow – Magenta* | 114.198 | 730.319 | < 0.05 |
|  | *Post – reversal* | *Yellow – Magenta* | −1007.544 | −379.356 | < 0.001 |
| B) $y = \beta_0 + \beta_{cluster}Cluster + \beta_{Age}Age$ | | | | | |
| | | $\beta$ | SE | t(df) | P |
| Vineland – |  |  |  |  |  |
| | Talking | $\beta_{cluster} = -2.04$ | 0.91 | −2.24 (33) | < 0.05 |
| | Caring | $\beta_{cluster} = -2.25$ | 1.03 | −2.19 (33) | < 0.05 |
| | Relating | $\beta_{Age} = 1.11$ | 0.42 | 2.63 (33) | < 0.05 |
| | Playing | $\beta_{cluster} = -2.26$ | 0.87 | −2.62 (33) | < 0.05 |
| | Large Muscles | $\beta_{Age} = 2.41$ | 0.53 | 4.54 (33) | < 0.001 |
| | Small Muscles | $\beta_{Age} = 1.31$ | 0.45 | 2.91 (33) | < 0.01 |
| | Total | $\beta_{cluster} = -11.85$ | 4.32 | −2.74 (33) | < 0.01 |
| | | $\beta_{Age} = 7.24$ | 1.89 | 3.82 (33) | < 0.001 |
| IBQ – |  |  |  |  |  |
| | Surgency | $\beta_{cluster} = -7.51$ | 3.20 | −2.34 (34) | < 0.05 |

There were missing data points in the questionnaires as some participants did not complete them. This left us with 36 participants on the Vineland (*N*_*Magenta*_ = 23, *N*_*Yellow*_ = 13) and 37 participants on the IBQ (*N*_*Magenta*_ = 24, *N*_*Yellow*_ = 13) questionnaires. Carrying out linear regression on questionnaire scores where the independent variables were cluster identity and age (months) revealed significant differences between clusters in terms of questionnaire scores and significant relationships between age and questionnaires. **Table 3B** and **Figure 6B** show decreased questionnaire scores for the yellow compared to the magenta cluster in terms of Vineland – Talking, – Caring for Self, – Playing and Using Leisure Time, – Total and IBQ – Surgency. There was a positive relationship between age and Vineland – Relating to Others, – Using Large Muscles, – Using Small Muscles, and Total (see **Table 3B**). There were no significant differences between the parents of infants who were assigned to different clusters in terms of STAI.

A higher variance ratio criterion (VRC) suggests a better separation between data clusters. In the next analysis, we clustered infants based on two separate sets of measures and compared the two clustering solutions on a third set of measures in terms of VRC. This involved: clustering solutions derived from i) behavioural measures and parameters, tested on questionnaire data and ii) questionnaire data and parameters, tested on behavioural measures. The behavioural measures involved four features: accuracy, perseveration, the difference between the dwell time on correct and incorrect ROIs in the pre-reversal phase and the post-reversal phase. The questionnaire data involved Vineland (7 features) and IBQ (3 features) subscales. Finally, the parameters involved the prior preferences over the animations *c* and the bias for the dominant policy (i.e., ROI side) ɛ. In this analysis, we included the infants who have questionnaire data (*n* = 36). Clustering solutions derived from parameters provided a better separation between clusters when tested on both the questionnaire data and behavioural measures (see **Table 4**).

**Table 4.** Comparisons between clustering solutions – The upper panel compares clustering solutions derived from behavioural measures and parameters when tested on the questionnaire data. The lower panel compares clustering solutions obtained with the questionnaires and parameters when tested on the behavioural measures. The entries in the table are the variance ratio criterion (VRC) estimates. Higher VRC indicates better separation across clusters.

| Test data: Questionnaires | Clustering |  |
| --- | --- | --- |
|  | Behavioural measures | Parameters |
|  | 0.95 | 2.85 |

**Table 4.** Comparisons between clustering solutions – The upper panel compares clustering solutions derived from behavioural measures and parameters when tested on the questionnaire data.
| Test data: Behavioural measures | Clustering |  |
| --- | --- | --- |
|  | Questionnaires | Parameters |
|  | 2.21 | 15.91 |

## Discussion

Here we demonstrate for the first time that is it is possible to reliably fit theory driven models to infant choices, to parameterise individual differences in behaviour at a mechanistic level. To do so, we employed a novel gaze-contingent version of a cued reversal paradigm, optimised to allow infants to act, with consequence, on the world; shaping their own learning experience in response to changes in the environment.

By choosing correctly or incorrectly in the decision-making period, infants subsequently saw a clear and blurred version of a cartoon clip, serving as rewarding and aversive stimuli respectively. In these two clips, visual (e.g. colour, luminance, velocity) and auditory features are matched, but the auditory-visual information differs in its coherence or value across the conditions. In this study, and in earlier pilots, we showed longer dwell times on the clear, relative to the blurred, animations suggesting infants find these stimuli more engaging. Similarly in adults it has been shown that people make fewer fixations and tend to have reduced viewing time on noisy versions of images compared to their original^49^. Optimising rewarding and punishing outcomes for infant studies requires careful consideration. The absence of reward e.g. not playing the animation at all, might seem like the obvious choice in a developmental context when common primary (e.g. electric shocks) and secondary (e.g. monetary losses) negative reinforcers are not appropriate. However, infants <12 months old will quickly lose interest when nothing is presented on the screen. As such, any peripheral measures such as EEG, NIRS or pupillometry will be confounded by infants looking away on the non-rewarding trial types. Additionally, commonly reported experimenter efforts to recapture attention (e.g. shaking a rattle or blowing bubbles) might artificially create high salience conditions which would confound the measure of key variables such as prediction errors. Here we believe that our paradigm – fully gaze-contingent, self-paced, with outcomes matched on as many features as possible – strikes the optimal balance between these considerations and might inform more tightly controlled paradigm design for studying infant learning in future.

At the group-level, our model agnostic results show reduced accuracy and increased perseveration in the post-reversal phase. There was also a decrease in the dwell time on the displayed animation following the reversal. This general tendency towards perseveration is widely reported in infant studies involving reversals^38,52,53^. In Piaget’s classic A-not-B paradigm, a toy is hidden initially at location A, and the infants explore and recover the toy at location A. Once the infants retrieve the toy successfully on a few trials, the toy is hidden at location B next time as the infant observes. It is widely reported that infants under 12 months old display perseverative errors by exploring location A rather than B after the location switch. Importantly, the perseverative errors in our cued-reversal task fit well with the previously reported infant behaviour on paradigms involving reversals, validating our novel gaze-contingent task against classic in-lab games and structured play paradigms that have historically been used to capture cognitive flexibility at these ages.

Crucially, however, we then fit these choices with a Markov decision process (MDP) formulation of active inference and estimate model parameters specific to each infant. Individual-level behaviour on the task can best be explained by a model comprising two parameters *c* and ɛ - which represent reward *preference or sensitivity,* and *dominant policy bias* (for the left or right of the screen). Clustering based on model parameter estimates yielded two clusters: magenta and yellow. The yellow cluster had an increased bias and decreased reward preference, compared to the magenta cluster. These parametric differences manifest in terms of distinct behavioural profiles in the pre- and post-reversal phases, with the yellow cluster showing greater accuracy followed by a larger drop following the switch, and more perseveration. In effect, this group demonstrate a ‘primacy bias’ in learning, such that they perform better until the reversal occurs, after which point they learn less, and seem relatively insensitive to the continued negative feedback. In contrast, the magenta cluster displayed a more exploratory decision-making profile, with decreased, though still above chance, accuracy pre-reversal and increased accuracy post-reversal. These infants might be said to show greater cognitive flexibility.

The yellow and magenta clusters did not differ in terms of age. Prior work on the development of perseveration in early infancy is mixed, with recent studies employing variants of the A-not-B paradigm suggesting that older infants might show more perseveration than their younger counterparts^54^. In the current study, however, these clusters show differences in terms of adaptive skills and temperament, as measured via established developmental assessments. The magenta cluster, with the more exploratory or flexible cognitive profile, generally showed better adaptive functioning assessed with the Vineland II ABS. This demonstrates the computational mechanisms supporting good cognitive flexibility also underscore adaptive behaviour from the very earliest stages of development. Furthermore, increased IBQ-R VSF surgency scores in the magenta cluster suggest increased extraversion compared to the yellow cluster. This might partially explain why the magenta cluster exhibits more exploratory behaviour in the cued reversal task. Whilst the infants in this study comprise a typically developing sample, and none of the participants would be considered ‘at-risk’ or high-likelihood of developmental delay, it is nonetheless important to demonstrate that robust parameterisations of infant behaviour can identify subgroups with distinct cognitive profiles that are otherwise ‘hidden’ in the behavioural data alone. Furthermore, we take the important step of demonstrating that clustering based on model parameters yields better solutions than clusters derived from behavioural measures and questionnaires when tested on unseen data, demonstrating the generalizability of the model-based solutions.

Our model was subject to rigorous validation, including model comparisons that can reveal information about the model architecture that infants are using to perform this task. For example, the first stage of model-comparisons revealed that infants do not carry forward their learned contingencies across blocks. There are several reasons why this might be the case, including a decreased learning rate or forgetting the contingencies during the short break between blocks. Indeed, working memory capacity has long been recognised as a pre-cursor to executive functions that contributes to performance on cognitive flexibility tasks in early life^55^. Additionally, at the second stage of model-comparisons, the model containing the novelty component of the expected free energy was not well supported by the model evidence. For an infant who makes correct decisions by sampling the left ROI in the pre-reversal phase, the novelty component would encourage sampling the right (incorrect) ROI simply to learn about the consequences of doing so^41^. The exclusion of the novelty component in the winning model suggests that, in our sample, behaviour is not yet characterised by this strategy.

The final stage of the model comparison yielded low confidence in the volatility *ω* and learning rate *η* parameters expressed in terms of posterior probabilities. These two parameters were fixed to their group-level estimates, and the prior preferences over the animations *c* and the bias for the dominantly sampled ROI ɛ were estimated for each infant. Fixing the learning rate *η* meant that the model still learned over trials, but this parameter no longer expressed individual differences between infants. A similar logic applies to the volatility *ω* parameter. Given, that computational indices of volatility and learning rate have proven to be influential in characterising neuropsychiatric differences in adult samples^10,14,19,22,56^, and response to pharmacological treatment^27,30,57^ it will be interesting for future longitudinal studies to consider at what point along the developmental trajectory the model architecture changes such that individual expressions of volatility and learning rate capture meaningful differences between individuals. This might represent an important window of opportunity for intervention. Volatility represents the rate of change in a contingency mapping across time^10,16,58^. Given that the infants in our study can only focus for ∼18 trials, with a single contingency reversal, it makes sense that volatility estimates don’t yet factor in to their individual decision-making strategies. Sustained attention, to be able to integrate over a wider envelope of time, is likely necessary for meaningful estimates of volatility. We stress, however, that we *could* have fit volatility *ω* and learning rate *η* parameters to our data, but model comparisons indicate they likely would not offer true or meaningful explanations of infant behaviour. As we see the early emerging steps towards infant behavioural modelling^59,60^, we urge for models fit to infant data to be held to the same standards as adults^43^ to arbitrate parsimoniously between competing explanations and the developmental emergence of cognitive processes.

Finally, we explored the relationship between pupil size and model-derived prediction errors as a measure of surprise. Comparing the post and pre-reversal responses yielded significant differences in pupil size and prediction error for *n* = 3. Moreover, these differences (pupil size *Pupil*_*Post*−*Pre*_ and prediction error *PE*_*Post*−*Pre*_) were significantly correlated for *n* = 3. Furthermore, pupil size and prediction errors were larger on incorrect trials across the task and the differences between incorrect and correct trials for pupil size *Pupil*_*Inc*−*Cor*_ and prediction error *PE*_*Inc*−*Cor*_ was also significantly correlated. These findings demonstrate that in infants, as in adults, pupillary responses signal surprise^14,16,27,28,61,62^. There is also evidence that noradrenergic (NA) activity, especially originating from Locus Coeruleus (LC), mediates pupil dilation^24,25,27^, and the interactions between cholinergic and noradrenergic systems are suggested to be involved in determining contextual changes^63,64^. Our work suggests that in infants, pupil dilations signal contextual switches where large prediction errors are to be expected. Future imaging studies – especially cutting-edge tools like high-density diffuse optical tomography (hdDOT^65–67^), could uncover the intricate relationship the cortical afferents which receive projections from the LC, such as the frontal and medial prefrontal cortex and their role in processing prediction errors.

This study showed that computational modelling can be a valuable method to distinguish between behaviourally distinct groups of infants, and it provides insight into the computational underpinnings of infant behaviour. The winning model explained the behaviour of the infants well, with a median decision fit of ≈ 86%, and also showed good parameter recoverability, identifiability and faithful simulations of behavioural performance. These results set the standards, show proof of concept, and demonstrate the potential, for the field of infant computational psychiatry. In particular, it will be important to discover to what extent these subgroups reflect either developmental stages or lasting cognitive subtypes, given the association of a ‘primacy bias’ in inference to rigid beliefs in adults^20^.

## Acknowledgements

R.P.L is supported by a Wellcome Trust Royal Society Henry Dale Fellowhip, and funding for these studies was provided by a Lister Institute Biomedical Research Prize and also a Wellcome Leap grant awarded to R.P.L. R.A.A. is a Future Leaders Fellow (MR/W011751/1). We would like to thank all the infants and their parents who gave up their time to take part in this study, and also Berk Mirza for all of his hard work in supporting the analysis.

