## Supplemental Materials for "Computational neurodevelopment: infant decision-making in changing environments"

### Hypothesis testing

We used generalised linear mixed-effects models (GLMMs) for hypothesis testing involving the response variables accuracy, perseveration, dwell time – decision period and dwell time – animation period. Table S1 shows the target distributions and link functions used in the GLMMs.

| Response variable | Target distribution | Link function |
|---|---|---|
| Accuracy | Binomial | Logit |
| Perseveration | Binomial | Logit |
| Dwell time (decision period) | Normal | Identity |
| Dwell time (animation period) | Normal | Identity |

**Table 1.** Target distributions and the link functions used in the GLMM analysis.

The fixed effects involved in the GLMM were gender, age, block (1$^{st}$ or 2$^{nd}$), reversal (pre or post-reversal) and the interaction between block and reversal. Additionally, we included the fixed effect of accuracy (correct or incorrect response) for the response variable *dwell time – animation period*.

The fixed effects involved in the GLMM analysis with the infant clusters included gender, age, block, reversal, cluster identity and the interactions between block and cluster identity, and reversal and cluster identity. The analysis involving the *dwell time – animation period* also included the interaction between accuracy and group.

The random effects in both analyses were block and reversal. We fitted models including and excluding each random effect and evaluated the evidence for models in terms of the Akaike information criterion (AIC). The reported statistics are those of the winning models (i.e., those with the least AIC).

### Computational modelling

#### Active inference and Markov decision process (MDP) models

In the active inference framework, the minimisation of Variational free energy underwrites perception, attention, and action. Variational free energy is expressed as

$$
\begin{aligned}
F &= -E_{Q(x)}[\ln P(\tilde{o},x)] - H[Q(x)] \\
&= -\ln P(\tilde{o}) + D_{KL}[Q(x) \,||\, P(x|\tilde{o})] \\
&= -\underbrace{E_{Q(x)}[\ln P(\tilde{o}|x)]}_{Accuracy} + \underbrace{D_{KL}[Q(x) \,||\, P(x)\,]}_{Complexity}
\end{aligned}
$$

Where $x$ represents the hidden variables, $\tilde{o}$ is the outcomes over time $\tilde{o} = (o_1, \dots, o_T)$, $P(\tilde{o}, x)$ is generative model of outcomes, $Q(x)$ is the approximate posterior distribution, $P(x|\tilde{o})$ is the true posterior distribution, $P(x)$ is the prior distribution, $H$ represents entropy, and $D_{KL}$ is the Kullback-Leibler (KL) divergence.

The first line in the equation above expresses free energy in terms of a generative model $P(\tilde{o}, x)$ and approximate posterior distribution $Q(x)$. The second line shows that the free energy is an upper bound on negative log model evidence $-\ln P(\tilde{o})$, as the KL divergence can never be less than zero. Minimising the KL-divergence makes the distribution $Q(x)$ an approximation of the true posterior distribution over the hidden variables $P(x|\tilde{o})$. The final line expresses free energy in terms of accuracy and complexity. Accuracy describes how accurately the outcomes can be explained under the (approximate) posterior beliefs about the hidden variables, while complexity describes the dissimilarity between the approximate posterior and the prior distribution over the hidden variables. Minimising free energy maximises accuracy while minimising complexity.

Markov decision process (MDP) models use discrete hidden states $s_t$ and time to describe how outcomes $o_t$ are generated at each time point. The states $s_t$ represent the hidden aspects of an environment (e.g., context), and the model has control over some of the states (e.g., gaze location) through policies $\pi$, where policies specify potential actions to pursue in the future. The model uses likelihood matrices $\mathbf{A}$ to express the probabilistic mapping from states to outcomes and transition matrices $\mathbf{B}$ to describe the probabilistic transitions between states. The model's prior beliefs over the outcomes and initial states are described in the $\mathbf{C}$ and $\mathbf{D}$ vectors. In this work, the hidden variables involve hidden states $\tilde{s} = (s_1, \dots, s_T)$ and policies $\pi$ and the model parameters that describe state-to-outcome mappings $\mathbf{A}$, where $P(o_t|s_t) = Cat(\mathbf{A})$. The model has approximate posteriors over these hidden variables and model parameters $Q(x) = Q(\tilde{s}, \pi, \mathbf{A})$. Here, we considered a hierarchical MDP, where the higher level defines the initial states at the lower level and for each time point of the higher level, the lower level has multiple state transitions. Figure S1 provides a generic graphical representation of hierarchical generative models. See Friston et al., 2017[1] and Heins et al., 2020[2] for more details.

The policies are evaluated in terms of the free energy expected in the future, namely expected free energy $G_\pi$. Crucially, active inference agents are equipped with the prior belief that the likely policies are those with low expected free energy. The prior beliefs about policies $P(\pi)$ and expected free energy are expressed as

$$P(\pi) = \sigma(-G_\pi)$$
$$G_\pi = \sum_{\tau>t} G_{\pi\tau}$$

Where $G_{\pi\tau}$ refers to the expected free energy evaluated at future time points $\tau > t$.

$$\begin{aligned} G_{\pi\tau} &= E_{Q(\mathbf{A}, o_\tau, s_\tau|\pi)}[\ln Q(\mathbf{A}, s_\tau|\pi) - \ln P(\mathbf{A}, o_\tau, s_\tau)] \\ &\approx -\underbrace{E_{Q(o_\tau, s_\tau|\pi)}\big[D_{KL}[Q(\mathbf{A}|o_\tau, s_\tau) \,||\, Q(\mathbf{A})]\big]}_{Novelty} \\ &\quad -\underbrace{E_{Q(o_\tau|\pi)}\big[D_{KL}[Q(s_\tau|o_\tau, \pi) \,||\, Q(s_\tau|\pi)]\big]}_{Epistemic\ vaue} - \underbrace{E_{Q(o_\tau|\pi)}[\ln P(o_\tau)]}_{Extrinsic\ value} \end{aligned}$$

Where $Q(\mathbf{A}, o_\tau, s_\tau|\pi) = Q(\mathbf{A}|o_\tau, s_\tau)Q(o_\tau, s_\tau|\pi)$ and $P(o_\tau) = Cat(\boldsymbol{C})$.

The expected free energy consists of three terms, namely *novelty*, *epistemic value*, and *extrinsic value*. *Novelty* and *epistemic value* underwrite uncertainty resolving behaviour. While the *novelty* component resolves uncertainty about the model parameters (e.g., mapping from states to the outcomes), *epistemic value* resolves uncertainty about the hidden states. *Extrinsic value* drives goal-oriented behaviour such that the policies that are likely to fulfil the model's preferences become more likely to be pursued[3].

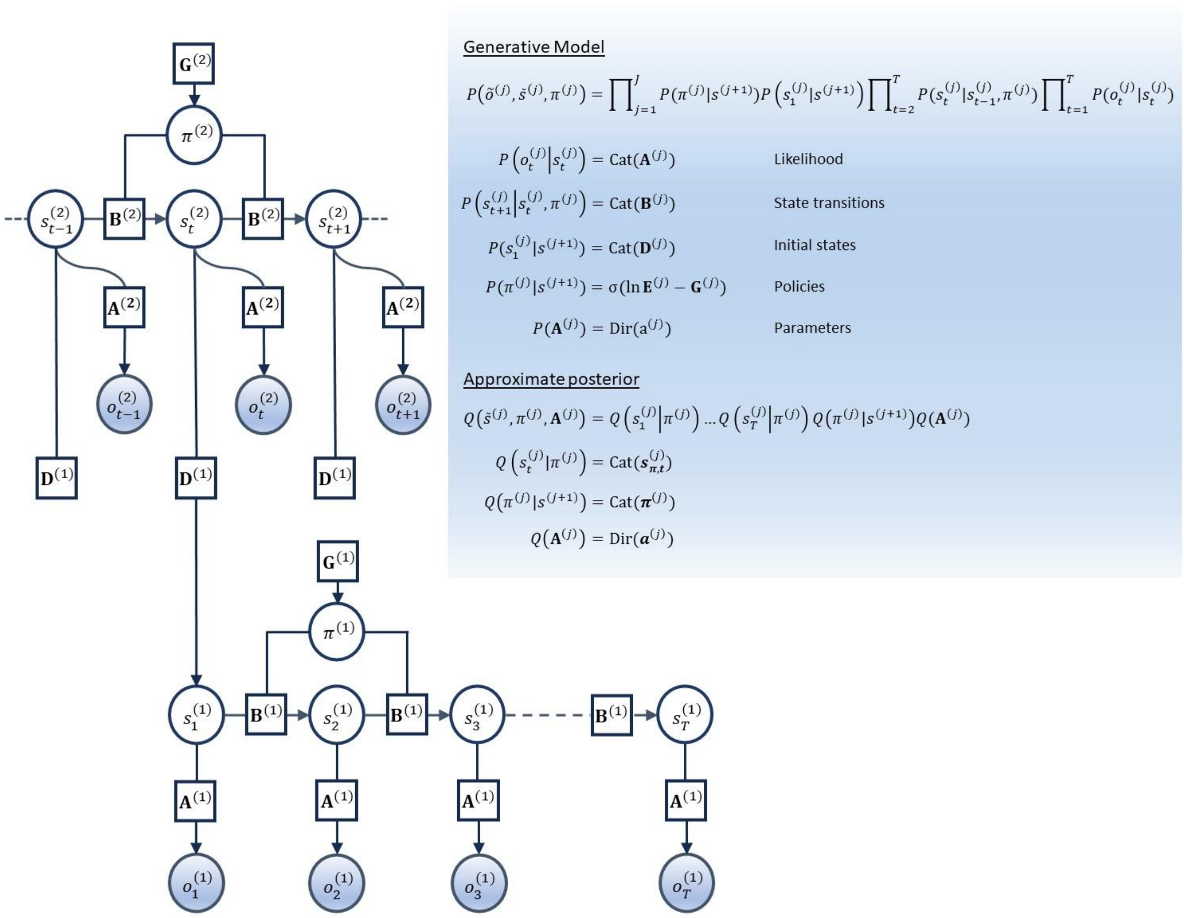


**Figure S1.** Generative model, priors and approximate posterior – The panel on the left describes the dependencies between different units in this hierarchical model in a similar fashion as in Friston (2017)[1]. Here, the squares represent model parameters, and white and blue circles represent the random variables and observable outcomes, respectively. The panel on the right describes the generative model, the priors and the approximate posterior over the hidden variables and model parameters. The superscript $j$ refers to the level of the hierarchy, whereas the subscripts refer to the time points. The bold letters correspond to the sufficient statistics of categorical and Dirichlet distributions. The prior likelihood concentration parameters are expressed in normal font $a$ to distinguish them from the posterior likelihood concentration parameters $\boldsymbol{a}$.

Inferring hidden states and policies entail minimisation of free energy with respect to the sufficient statistics of the posterior distributions over the respective variables. Below equations express the model's beliefs about the hidden states:

$$\boldsymbol{s}_{\boldsymbol{\pi},\boldsymbol{\tau}=\mathbf{1}}^{(j)} = \sigma\left(\underbrace{\ln \mathbf{D}^{(j)}\, \boldsymbol{s}_{\boldsymbol{\tau}}^{(j+1)}}_{\substack{Priors\ about \\ initial\ states}} + \underbrace{\omega \ln \mathbf{B}_{\boldsymbol{\pi},\boldsymbol{\tau}}^{(j)} \cdot \boldsymbol{s}_{\boldsymbol{\pi},\boldsymbol{\tau}+\mathbf{1}}^{(j)}}_{Backward\ message} + \underbrace{\ln \mathbf{A}^{(j)} \cdot o_{\tau}^{(j)}}_{Evidence} + \underbrace{\ln \mathbf{D}^{(j-1)} \cdot \boldsymbol{s}_{\mathbf{1}}^{(j-1)}}_{\substack{Evidence\ from \\ lower\ level}}\right)$$

$$\boldsymbol{s}_{\boldsymbol{\pi},\boldsymbol{\tau}>\mathbf{1}}^{(j)} = \sigma\left(\underbrace{\omega \ln \mathbf{B}_{\boldsymbol{\pi},\tau-1}^{(j)}\, \boldsymbol{s}_{\boldsymbol{\pi},\boldsymbol{\tau}-\mathbf{1}}^{(j)}}_{Forward\ message} + \underbrace{\omega \ln \mathbf{B}_{\boldsymbol{\pi},\boldsymbol{\tau}}^{(j)} \cdot \boldsymbol{s}_{\boldsymbol{\pi},\boldsymbol{\tau}+\mathbf{1}}^{(j)}}_{Backward\ message} + \underbrace{\ln \mathbf{A}^{(j)} \cdot o_{\tau}^{(j)}}_{Evidence} + \underbrace{\ln \mathbf{D}^{(j-1)} \cdot \boldsymbol{s}_{\mathbf{1}}^{(j-1)}}_{\substack{Evidence\ from \\ lower\ level}}\right)$$

Here, $\tau$ corresponds to epochs representing time points in the past, present and future. For example, after seeing the star cue in the cued-reversal paradigm at $t = 1$, the model forms expectations about the animations at a future epoch $\tau = 2$. The inferences about the states depend on four terms. At the initial epoch $\tau = 1$, the expectations about the states at the higher level $\boldsymbol{s}_{\boldsymbol{\tau}}^{(j+1)}$ enter the lower level through the $\mathbf{D}$ matrix, which expresses the conditional probability of the initial states at the lower level, given the states at the higher level. For epochs $\tau > 1$, the first term is replaced by the forward messages. The forward and backward messages propagate posterior beliefs about states in the previous $\boldsymbol{s}_{\boldsymbol{\pi},\boldsymbol{\tau}-\mathbf{1}}^{(j)}$ and future epochs $\boldsymbol{s}_{\boldsymbol{\pi},\boldsymbol{\tau}+\mathbf{1}}^{(j)}$ to the current epoch under the model's beliefs about state transitions $\mathbf{B}$. The evidence term expresses the likelihood of the states yielding the observed outcomes through the $\mathbf{A}$ matrix. The ultimate term is an ascending message from the level below $\boldsymbol{s}_{\mathbf{1}}^{(j-1)}$, and it supplies evidence for the states at the current level of the hierarchy after cycling through the time points at the lower level.

Ultimately, the forward and backward messages are responsible for integrating evidence obtained over different epochs on a given level of the hierarchy. The parameter $\omega$ expresses the extent to which the states in the previous $\boldsymbol{s}_{\boldsymbol{\pi},\boldsymbol{\tau}-\mathbf{1}}^{(j)}$ and future epochs $\boldsymbol{s}_{\boldsymbol{\pi},\boldsymbol{\tau}+\mathbf{1}}^{(j)}$ influence the beliefs in the current epoch via the forward and backward messages. This is a precision parameter, and it defines the model's beliefs about the state transitions as below:

$$P(s_{\tau+1} = m | s_{\tau} = n, \pi, \omega) = \frac{\overline{\mathbf{B}}_{mn}^{\omega}}{\sum_m \overline{\mathbf{B}}_{mn}^{\omega}}$$

In the context of the cued reversal paradigm, $\overline{\mathbf{B}}$ represents an identity matrix expressing the transitions between contexts (i.e., pre- and post-reversal phases). The higher the $\omega$ parameter, the more the evidence from the previous and next epochs can be integrated with the beliefs at the current epoch. When $\omega$ is very high, the model would believe that the transitions between contexts are not likely, whereas with very low $\omega$, the model would expect a volatile environment (i.e., frequent changes between contexts).

The posterior beliefs about the policies is defined as a softmax function of three terms, namely bias in policies $\mathbf{E}$, free energy of policies $\mathbf{F}$ and expected free energy $\mathbf{G}$.

$$\boldsymbol{\pi} = \sigma(\ln \mathbf{E} - \mathbf{F} - \mathbf{G})$$

Learning model parameters entails updating model parameters to minimise free energy. The parameters of the likelihood matrix are defined in terms of the concentration parameters of Dirichlet distribution. These parameters are updated as below:

$$\boldsymbol{a} = a + \eta \sum_{\tau} \boldsymbol{s_\tau} \otimes o_\tau$$

$$\boldsymbol{s_\tau} = \sum_{\pi} \boldsymbol{s_{\pi,\tau}}\, \boldsymbol{\pi}$$

Where $a$ and $\boldsymbol{a}$ are the prior and posterior concentration parameters, respectively, $\boldsymbol{s_\tau}$ is the Bayesian model average of the beliefs about the states given policies $\boldsymbol{s_{\pi,\tau}}$ with the posterior over the policies $\boldsymbol{\pi}$, $o_\tau$ is the outcome vector with a probability of 1 is assigned to the observed outcome and $\eta$ is the learning rate.

The posterior concentration parameters are converted to probabilities to obtain the new likelihood matrix

$$E_{P(\mathbf{A}|\boldsymbol{a})}[\mathbf{A}_{ij}] = \frac{\boldsymbol{a}_{ij}}{\sum_i \boldsymbol{a}_{ij}}$$

#### Model of the cued reversal paradigm

The MDP model of this paradigm consisted of a hierarchy with two levels. The higher level of the model involved a single state factor, and outcome modality labelled *context*, which encoded the pre- and post-reversal phases in the paradigm. There was an identity mapping from the context states to the context outcomes on this level.

The lower level of the model involved three state factors labelled *context*, *starting side*, *when*. The levels of the context states were the same as at the higher level, and the prior beliefs about the initial states at the lower level were defined as conditional probabilities as they depended on the states at the higher level $P(s_1^{(1)}|s^{(2)})$. The *starting side* states had two levels corresponding to the infants' decision on the first trial, either the *left* or the *right* ROI. The *starting side* and *context state* factors determined the ROI where the clear and blurred animations are displayed, either left or right. The *when* state factor had three levels encoding time and chosen actions in the decision-making period. These were labelled i) $t = 1$, ii) $t = 2, Left\ chosen$, and iii) $t = 2, Right\ chosen$. At $t = 1$, the star cues were presented and depending on the chosen action (left or right), either a clear or blurred version of the animation was displayed at $t = 2$. The states $Left\ chosen$ and $Right\ chosen$ described the chosen ROI in the decision period. There was a single outcome modality representing the *visual* outcomes. The *visual* outcomes had four levels, namely the orange and blue star cues and the clear and blurred versions of the animation.

The outcomes were generated in a way that the first level of the when states, $t = 1$, mapped onto the orange and blue star cues in the pre- and post-reversal phases, respectively. Given that the *starting side* is left, the $t = 2, Left\ chosen$ and $t = 2, Right\ chosen$ levels of the *when* states mapped onto the clear and blurred animations in the pre-reversal phase. The same states mapped onto the blurred and clear animations in the post-reversal phase. The mappings in the pre- and post-reversal phases were swapped for when the *starting side* is right compared to when it is left (see Figure S2 left panel).

We considered two kinds of generative models: i) a model with four matrices, including two context states for each level of the *starting side* factor (*left* and *right*), and ii) a model with two matrices, including two context states only. In both models, we defined the prior expectations about the context states at the higher level such that the first context state was much more likely. Initially, the model does not know the state–outcome mappings; thus, the context states represent uniform beliefs over the outcomes. The contexts are sculpted via learned likelihood mappings as the model performs the task. This motivated our choice of prior beliefs, meaning there should be no reason to expect multiple contexts to be likely at the beginning when contexts do not represent concrete structures expressed in terms of likelihood mappings. Practically, this is implemented by assigning the states the following concentration parameters $\mathbf{d}^{(2)} = [1 \quad 0.01]$, where $P\left(s_1^{(2)}\right) = Cat(\mathbf{D}^{(2)})$ and $\mathbf{D}^{(2)} \sim dir(\mathbf{d}^{(2)})$. Essentially, with these beliefs, the model would mainly update the concentration parameters of the likelihood mapping associated with the first context state in the lower level until the posterior beliefs for the second context state overtake. This is likely to happen when the first context state can no longer explain the model's observations; thus, the second context state would become more probable.

For an infant who chose opposite sides on the first trial of each block, there need to be four likelihood matrices to learn the true state–outcome mappings (see Figure S2A). The first model has the capacity to learn the likelihood mappings for all unique combinations of *context* and *starting side* states (see Figure S2B left panel). In this model, we set the prior probability of the *starting side* state that matches the infant's choice on the first trial to 1. With these prior beliefs the model would only update the concentration parameters of the likelihood mapping associated with the appropriate level of the *starting side* factor.

$$P\left(s_1^{(1),2}\right) = \begin{cases} [\ 1 \quad 0\ ]^T, & \text{First choice: Left} \\ [\ 0 \quad 1\ ]^T, & \text{First choice: Right} \end{cases}$$

where the bracketed and unbracketed numbers in the superscript correspond to the first level in the hierarchy and the *starting side* (2nd) state factor, whereas the subscript represents the first time point. *First choice* is the infant's choice on the first trial of a block. The first and second entries in the vectors above represent the starting sides: *left* and *right*.

The second model excluded the *starting side* states and would have to overwrite its learned contingencies if different starting sides were chosen across blocks (see Figure S2B right panel). Both models were initialised with small and uniform concentration parameters over the outcomes that can be observed in their relative time points (i.e., orange and blue star cues at $t = 1$, clear and blurred animations at $t = 2$, see Figure S2B). We will refer to the models including and excluding the starting side states as *complex* and *simple* models. Here, *complex* is a reference to the extra state factor (starting side) in the model. For infants with second block data, the prior likelihood concentration parameters were set to the posterior concentration parameters at the end of the first block in both models.

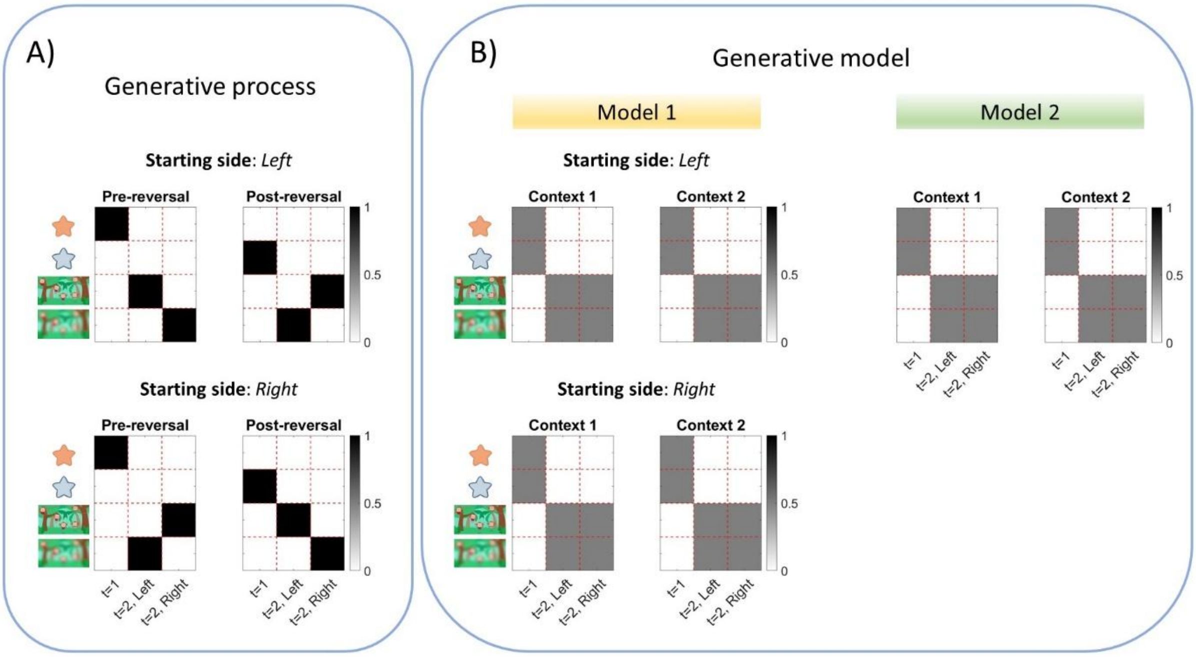


**Figure S2.** Generative process and model likelihood matrices – A) This panel shows the mapping from states (columns) to the outcomes (rows) in the generative process for different starting sides (left and right) and different phases of the paradigm (pre- and post-reversal phases). B) These panels show the prior likelihood mappings for two kinds of generative models. The first model has the same number of matrices as in the generative process, giving the model the capacity to learn distinct likelihood mappings associated with different starting sides and phases of the paradigm. The second model does not include the starting side as a state factor; therefore, there are only two matrices. The matrices in the generative process are labelled pre- and post-reversal. In contrast, the matrices in the generative models are labelled contexts 1 and 2. This is because the degree to which the learned contingencies are stored in the pre- and post-reversal phases depends on the model's capacity to recognise the change in contexts.

#### Model parameters

The parameters that were estimated on an individual basis by fitting the model to the data involved i) (inverse) volatility $\omega$, preferences $c$, learning rate $\eta$, and bias for choosing left or right ROI ε. The inverse volatility $\omega$ modulates the extent to which the expectations about the states in the previous $\boldsymbol{s}_{\boldsymbol{\pi},\boldsymbol{\tau}-\mathbf{1}}^{(j)}$ and future epochs $\boldsymbol{s}_{\boldsymbol{\pi},\boldsymbol{\tau}+\mathbf{1}}^{(j)}$ influence the states at the current epoch $\boldsymbol{s}_{\boldsymbol{\pi},\boldsymbol{\tau}}^{(j)}$. A very high (inverse) volatility $\omega$ equips the agent with the belief that the environment is very rigid and the switches between contexts are unlikely, whereas a low $\omega$ means that the contextual switches are to be expected frequently (see Figure S3A). The entries of the $C$ vector correspond to the visual outcomes in the cued reversal paradigm, and they define the model's preferences over the outcomes $P(o_\tau) = Cat(\mathbf{C})$. The entries in this vector are relative log probabilities (utilities), which are converted to probabilities using a softmax function. We defined the $C$ vector such that orange and blue star cues had no utilities, and the clear and blurred animations had utilities $c - 5$ and $5 - c$, respectively. Although we expected the infants to prefer the clear animation ($c > 5$), the formulation of the $C$ vector allowed preferences for the blurred animation as well ($c < 5$); see Figure S3B. The parameter $\eta$ modulated the rate at which the model learned the mapping from the states to the outcomes in terms of Dirichlet parameters. With a higher learning rate $\eta$, the model would update the state to outcome mappings to a larger extent (see Figure S3C). Finally, the predominantly chosen ROI was calculated for each infant and block individually, and the most and least frequently visited ROIs were assigned ε and $10 - \varepsilon$, respectively. These values were converted to probabilities by applying a softmax function to define the bias in policies (Left bias: $\mathbf{E} = \sigma([\ \varepsilon\ 10 - \varepsilon\ ])$, Right bias $\mathbf{E} = \sigma([\ 10 - \varepsilon\ \varepsilon\ ])$); see Figure S3D.

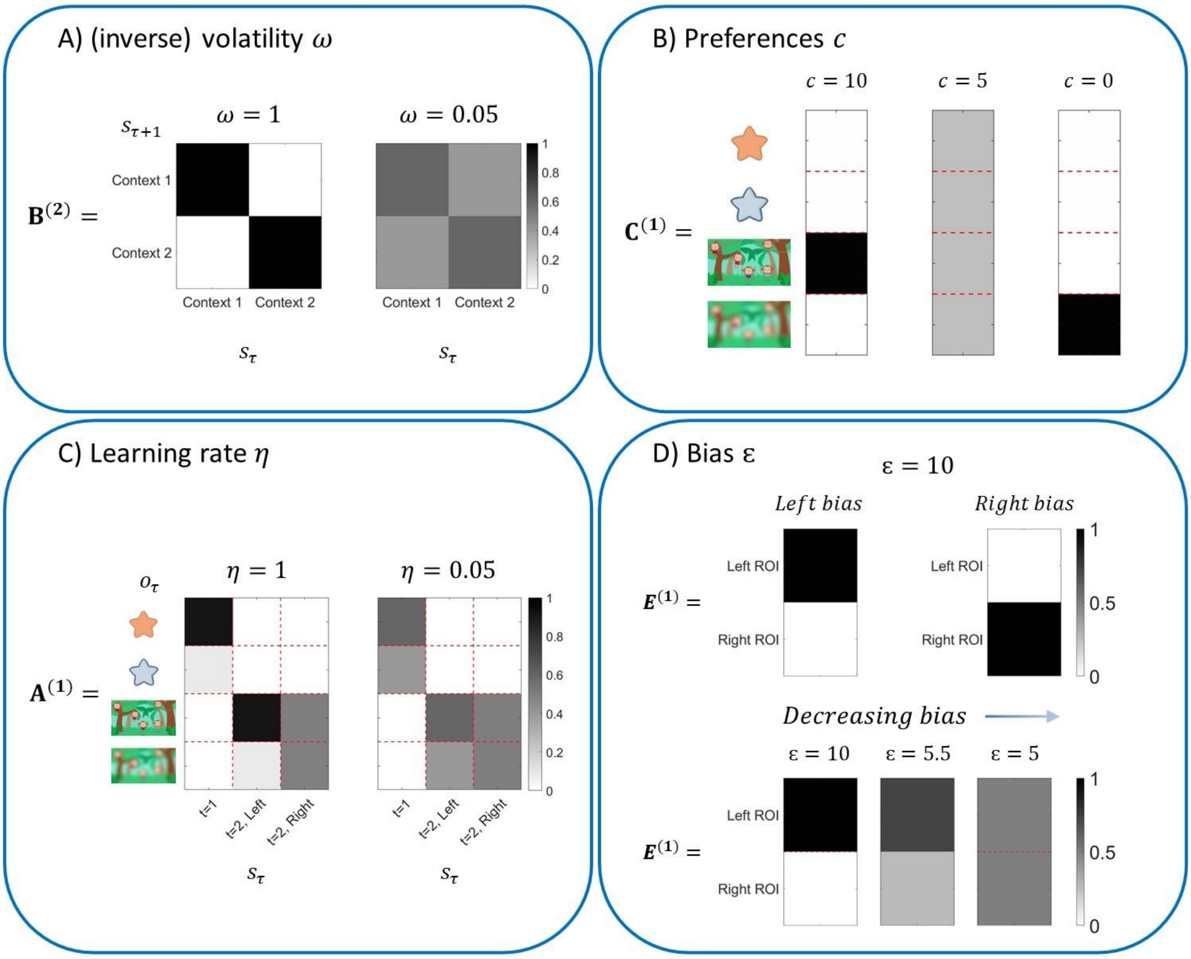


**Figure S3.** Parameters of the model – There are four parameters that are estimated per individual, namely (inverse) volatility $\omega$, preferences $c$, learning rate $\eta$, and bias ε. A) The transitions between context states are not likely across trials when $\omega = 1$. In contrast, the transitions between the context states become more likely when $\omega = 0.05$. B) The $c$ parameter modulates the model's preferences. The model prefers the clear animation when $c > 5$ (left vector), the blurred animation when $c < 5$ (right vector) and neither animation when $c = 5$ (centre). C) The model learns the likelihood matrices faster when the learning rate is $\eta = 1$ than when it is $\eta = 0.05$. D) The model has a strong bias for choosing the left and right ROIs when $\varepsilon = 10$. The ROI for which the infants are biased is estimated for each infant and block individually, and the predominantly chosen side is assigned the parameter ε, whereas the opposite ROI is assigned $10 - \varepsilon$. These entries are converted to probabilities with the softmax function. The panels at the bottom show that the bias for a given ROI (e.g., left) decreases as ε decreases from $10$ to $5$.

#### Model comparison

In the first stage of model comparison, we compared the *complex* model and three variations of the *simple* model in terms of free energy (evidence). In the first variation of the *simple* model, the prior likelihood concentration parameters at the beginning of the second block were set to the posterior likelihood concentration parameters at the end of the first block. We will refer to this model as *simple – keep experience* as the model carries its experiences from the first block forward to the second block. In the second variation, the likelihood mappings at the beginning of the second block were reset, and the learned mappings in the first block were discarded. We refer to this model as *simple – discard*. In the third variation, we defined the prior concentration parameters of the initial states at the higher level uniformly $\mathbf{d}^{(2)} = [1 \quad 1]$. This is a special case that prevents the model from learning separate likelihood mappings associated with the pre- and post- reversal phases. In other words, the model overwrites its experiences in the pre-reversal phase with the post-reversal phase and the likelihood matrices under each context state are always identical. We will refer to this model as the *simple – no context* since it cannot learn unique contexts.

In the second stage of model comparison, we fit models by excluding each component of the expected free energy – one at a time. *Novelty* is related to the curiosity fulfilling drives, as it encourages the agent to visit rarely occupied states to learn their mapping to the outcomes. In the cued reversal paradigm, a model without *novelty* would have no motive to resolve uncertainty about the likelihood concentration parameters (i.e., mapping from states to the outcomes). For example, the *novelty* component would encourage an agent that makes successive correct choices in the pre-reversal phase to sample the opposite (incorrect) ROI to learn the consequences of occupying that state. Excluding *epistemic value* robs the model of its drive to resolve uncertainty about the states. This component encourages the model to sample the ROIs that would resolve uncertainty about the context. A model without *extrinsic value* would have no preferences to fulfil. In the current paradigm, this translates to having no preferences over the clear and blurred versions of the animation.

The final stage of model comparison involved estimating evidence for models defined in terms of parameter combinations. This approach employs a general linear model consisting of a term encoding group averages per parameter in a hierarchical inversion scheme, namely parametric empirical Bayes (PEB). The evidence for models including and excluding parameters is estimated using Bayesian model reduction (BMR)[4]. This method allows the posterior probabilities of the model parameters to be estimated. The parameters with low posterior probabilities ($< 0.5$) were fixed to their posterior expectations at the group level, and the model inversion was repeated for each infant.

In brief, model comparison involved three stages where different models were compared in terms of their free energy. In the first stage, we compared four models: i) *simple – keep experience*, ii) *simple – discard*, iii) *simple – no context*, and iv) *complex*. In the second stage, the winning model of the first stage was refitted to the data, excluding each component of the expected free energy (EFE) and compared with the original EFE. These models were: i) *EFE*, ii) *EFE – without novelty*, iii) *EFE – without epistemic value*, and iv) *EFE – without extrinsic value*. In the third stage, models defined in terms of the combination of model parameters were evaluated using PEB and BMR. Because there were four free parameters, each of which can be either included or excluded to define a model, there were $2^4 = 16$ models to compare at this stage. The redundant parameters were fixed at their group-level means, and model inversion was repeated.

#### Prediction errors

Prediction errors express a mismatch between the model's expectations and its observations. Here, we define prediction errors on both levels of the model.

##### Lower-level prediction error

The prediction errors at the lower level express the divergence between the observed animation and the model's expectation about the clear and blurred versions of the animation. The model's expectations about the animations are obtained by

$$Q\left(o_{\tau=2}^{(1)}\right) = \sum_{s_{\tau=2}} P(o_{\tau=2}^{(1)}|s_{\tau=2}^{(1)}) \underbrace{\sum_{\pi} Q\left(s_{\tau=2}^{(1)}|\pi^{(1)}\right)Q(\pi^{(1)})}_{Q(s_{\tau=2}^{(1)})}$$

At the lower level, the model observes the animations at the second time point $t = 2$. Therefore we are interested in the model's expectation about the second epoch $\tau = 2$ after observing one of the star cues at $t = 1$. The posterior predictive density over the hidden states

$Q(s^{(1)}_{\tau=2})$ is obtained by taking the Bayesian model average of posterior beliefs about states given policies $Q\left(s^{(1)}_{\tau=2}|\pi^{(1)}\right)$ with the posterior beliefs about the policies $Q(\pi^{(1)})$. Multiplying the likelihood term $P(o^{(1)}_{\tau=2}|s^{(1)}_{\tau=2})$ with the expected states $Q(s^{(1)}_{\tau=2})$ and marginalising over the states yields the model's expectations about the animations $Q\left(o^{(1)}_{\tau=2}\right)$. The prediction error is expressed as a KL-divergence between a categorical distribution representing the observed animation $R(o^{(1)}_{t=2})$ and the model's expectation about the clear and blurred versions of the animation $Q\left(o^{(1)}_{\tau=2}\right)$.

$$PE^{(1)} = D_{KL}\left[\, R\left(o^{(1)}_{t=2}\right) \,||\, Q\left(o^{(1)}_{\tau=2}\right) \right]$$

Here, $PE^{(1)}$ represents the prediction error at the lower level. The observed clear animation is expressed by $R\left(o^{(1)}_{t=2}\right) = [\,1 \quad 0\,]^T$, and the blurred animation by $R\left(o^{(1)}_{t=2}\right) = [\,0 \quad 1\,]^T$. From the model's perspective, the only possible outcomes at the second time point were the clear and blurred animations. Therefore, we removed the entries associated with the star cues when deriving the expectations about the outcomes $Q\left(o^{(1)}_{\tau=2}\right)$.

##### Higher-level prediction error

At the higher level, the model's expectations about the context states after observing the animation on a given trial can be taken as its prior expectation about the context states on the next trial. The prediction error at the higher level is expressed as a KL-divergence between the model's expectations about the context states on consecutive time points (i.e., trials at this level). At the initial time point, this term is defined as a KL-divergence between the model's posterior beliefs about the context states at the end of the first trial $Q\left(s^{(2)}_1\right)$ and its prior beliefs about the initial context states $P\left(s^{(2)}_1\right) = Cat(\mathbf{D}^{(2)})$, where and $\mathbf{D}^{(2)} \sim dir(\mathbf{d}^{(2)})$ and $\mathbf{d}^{(2)} = [1 \quad 0.01]$. With these Dirichlet concentration parameters, the model's prior beliefs about context is $P\left(s^{(2)}_1\right) \approx [\,0.99 \quad 0.01\,]^T$.

$$PE^{(2)} = \begin{cases} D_{KL}\left[\, Q\left(s^{(2)}_1\right) \,||\, P\left(s^{(2)}_1\right)\right], & t = 1 \\ D_{KL}\left[\, Q\left(s^{(2)}_{t+1}\right) \,||\, Q\left(s^{(2)}_t\right)\right], & t > 1 \end{cases}$$

Here, $PE^{(2)}$ represents the prediction error at the higher level. The resulting prediction errors from both levels were converted to z-scores for each infant separately.

Considering that the prediction errors from different levels of the model contribute varying amounts to the infant's physiological response to surprise where pupil dilation was used as a proxy, we fit separate multilinear regression models involving the intercept term and a separate weight for the prediction errors to the normalised pupil data of each infant.

#### Clustering

We applied k-means clustering on the estimated parameters to test if the clustering solution finds distinct infant subgroups in terms of their behavioural responses and questionnaire scores. Each parameter was rescaled using minmax normalisation over infants, and cosine distance was used to evaluate cluster membership. Cosine distance is defined in terms of cosine similarity, where the cosine similarity expresses the similarity between two vectors. Cosine similarity ($CS$) is defined as

$$CS = \cos\theta = \frac{V_1 \cdot V_2}{\|V_1\|\|V_2\|}$$

$$CD = 1 - CS$$

In the first equation, the numerator is the dot product between two vectors and the denominator is a product between the L2-norm of each vector. The second equation describes cosine distance ($CD$) in terms of cosine similarity. Here, we used two normalised vectors i) one with entries corresponding to an infant's parameter estimates, and ii) another vector representing the cluster centre (centroid) in terms of the average over infants in the cluster. We tested clustering solutions that allowed up to 10 clusters in terms of Calinski-Harabasz variance ratio criterion (VRC) where a greater VRC suggests a better clustering solution[5]. VRC is defined is terms of a ratio of between and within cluster variance

$$VRC = \left(\frac{SS_{Between}}{k-1}\right) \Big/ \left(\frac{SS_{Within}}{n-k}\right)$$

where $SS_{Between}$ and $SS_{Within}$ are between and within cluster sum of squares, respectively, $k$ is the number of clusters and $n$ is the number of data points (infants).
